# Structural insights into the inactive state of the adhesion GPCR ADGRV1

**DOI:** 10.64898/2026.03.05.709805

**Authors:** Yasmine Achat, Marie S. Prevost, Ariel Mechaly, Mariano Genera, Baptiste Colcombet-Cazenave, Armel Bezault, Jean-Marie Winter, Catherine Venien-Bryan, Bertrand Raynal, Pierre Lafaye, Patrick England, Gabriel Ayme, Massimiliano Bonomi, Laurent Prezeau, Nicolas Wolff

## Abstract

Adhesion G protein-coupled receptors (aGPCRs) are involved in numerous physiological processes, including cell-cell and cell-matrix interactions, and are associated with several human diseases. ADGRV1 is a member of the aGPCR family and plays a significant role in the sensorineural systems. Mutations of ADGRV1 are linked to the Usher syndrome, a genetic disorder causing deafness and blindness in human. However, the molecular mechanisms that control the activity of ADGRV1 remain unclear. In this study, we present the high-resolution cryo-electron microscopy structure of the inactive ADGRV1 receptor in complex with the nanobody RE02, providing detailed insights into its transmembrane domain, intracellular loop conformations and the inactive orthosteric site. Functional cellular assays revealed that ADGRV1 exhibits a weak constitutive activation independent of the tethered agonist peptide, primarily coupling with G_i_ proteins. These findings suggest that ADGRV1 may employ an alternative activation mechanism, distinct that from other aGPCRs reported so far. Our structural analysis highlights that ADGRV1 does not follow the conventional tethered agonist activation mechanism. This is due to the divergent sequence of its tethered agonist peptide, as well as the lack of residues critical for conformational transitions toward the active state in other aGPCRs. Moreover, the large intracellular loop 3 (ICL3) adopts a closed conformation, tightly packed against the transmembrane region of ADGRV1. This suggests that the ICL3 loop can compete with G-protein binding, ultimately acting as an additional barrier for ADGRV1 activation.

**Significance Statement:** Adhesion G protein–coupled receptors (aGPCRs) regulate essential processes such as cell communication and sensory function, yet the mechanisms controlling many family members remain poorly understood. We report the first high-resolution cryo–electron microscopy structure of the human receptor ADGRV1, a protein linked to Usher syndrome, a major genetic cause of deafness and blindness. Structural and functional analyses reveal that ADGRV1 displays weak constitutive signaling and likely operates through an activation mechanism distinct from the canonical tethered agonist model that defines most aGPCRs. Unique structural features, including a closed intracellular loop (ICL3) that may limit G-protein engagement, suggest an alternative regulatory strategy. These findings expand the conceptual framework of aGPCR activation and provide a foundation for understanding ADGRV1 function in sensory physiology and disease.

## Introduction

Adhesion G protein-coupled receptors (aGPCRs) form the second-largest family of GPCRs (GPCR family B2) with 33 members in humans^1^. They play crucial roles in transducing mechanical forces and mediating cell-cell and cell-matrix interactions, serving as primary regulators of various physiological processes such as cell polarity, adhesion, migration and differentiation. Subsequently, aGPCRs are involved in many pathophysiologies, including cancer development or nervous system disorders in humans ^1^.

According to phylogenetic classification, ADGRV1 is the sole member of the IX sub-family of aGPCR ^2^. Over the years, it has been known by several names, including GPR98, MASS1, Neurepin and VLGR1. ADGRV1 is expressed in the nervous system, particularly in sensory neurons involved in hearing and vision, and in astrocytes. Mutations of *ADGRV1* are responsible for the sensory defects associated with Usher syndrome type IIC ^3^. This rare disease is the most common genetic cause of combined deafness, balance defect and progressive blindness, affecting the sensory cells of the inner ear and the retina. ADGRV1 is a component of interstereocilia links that are required for the normal development of hair bundles exposed at the surface of the sensory hair cells in the organ of Corti ^4^. In the retina of zebrafish, expression impairment of ADGRV1 results in an alteration of the circulation of photopigments such as rhodopsin in photoreceptors, leading to a drop in the visual capacity of these animals ^5^. ADGRV1 is also linked to epilepsy in humans, likely related to its expression in the developing nervous system ^6^, where it regulates cell migration of astrocytes and GABA-ergic interneurons in the auditory cortex ^7 8 9^. Proteomics experiments on ADGRV1 in eukaryotic model cells have allowed identification of cellular partners involved in processes such as Ca^2+^ homeostasis in mitochondria and autophagy ^10 11 12 13^. Therefore, although the scaffolding function of ADGRV1 has been well-documented for the proper development of hair cells, the role of the receptor in cellular signalling and homeostasis of cochlear cells or photoreceptors is not yet known.

ADGRV1 is the largest GPCR identified to date, comprising 6,306 amino acids in humans (UniprotKB N°Q8WXG9). Multiple isoforms of ADGRV1 have been described, with up to eight reported ^14^. The full-length protein is predominantly expressed in the cochlea and the retina. Its long extracellular N-terminal fragment (NTF) comprises 5,879 residues, featuring 39 predicted calcium-binding CalX-β domains, 7 EAR (Epilepsy-associated repeat) domains, one PTX (Pentraxin) domain, and one GAIN (GPCR autoproteolysis-inducing) domain that includes the GPCR autoproteolytic cleavage site (GPS). Downstream the canonical seven-transmembrane helix domain of GPCR (7TMD), ADGRV1 possesses a 148-residues long cytoplasmic domain of unknown function. The 7TMD and the cytoplasmic domain together constitute the C-terminal fragment of ADGRV1 called the CTF. Experimental evidence suggests that autocleavage of ADGRV1 at the GPS removes the extracellular NTF (Figure S1) and exposes a short sequence of roughly 15-25 residues at the N-terminus of CTF, known as the Stachel peptide. This segment could act as a tethered agonist peptide (TA), activating ADGRV1. It has been proposed that the autocleavage of ADGVR1 results in a switch in the G protein coupling from Gα_s_, which is constitutively coupled to the full-length, uncleaved ADGRV1, to Gα_i_-mediated signalling by the cleaved ADGRV1, which is called ADGRV1β ^15 16 10^. However, seemingly contradictory experimental data have been published regarding the role of the Stachel peptide in ADGRV1 activation^17^. Moreover, the molecular mechanisms of ADGRV1 activation still remain unknown, although they have been reported for several other aGPCRs over the past three years, primarily through the structural characterization of their active forms by cryo-electron microscopy (Cryo-EM) ^18 19 20 21 22 23 24 25^.

Here, we determined the high-resolution structure of ADGRV1β in its inactive state by cryo-EM, in complex with a nanobody designed to stabilize the receptor. Structural comparison with other aGPCRs allows us to propose hypotheses to explain the weak G protein coupling observed in cells and the lack of activation by the Stachel peptide under our experimental conditions.

## Results

ADGRV1 is the largest receptor within aGPCRs and contains 5,891 residues in its extracellular domain, making it impractical to study the complete protein *in vitro*. In this study, we focused on producing and structurally characterizing the C-terminal fragment (CTF) of ADGRV1, referred to as ADGRV1β. This fragment starts at the unveiled N-terminus Stachel peptide (a short sequence of 15 residues between the GPS and the first transmembrane helix TM1) and includes the transmembrane domain (TMD) of 252 residues and the intracellular C-terminal domain (ICD) of 148 residues (**Figure S1**). ADGRV1β has been reported to be functional in cells, mediating signalling through the heterotrimeric Gi protein ^15^ and we have recently shown that it localizes in mouse hair cells up to postnatal day 21 ^26^.

### ADGRV1β and G_i_ signalling

We first determined the G protein signalling abilities of wild-type ADGRV1β (WT, of sequence -SVYAVYA-) using Bioluminescence Resonance Energy Transfer BRET technology in cellular assays. For this purpose, cells were transfected with the Gα-RLuc and βγ-Venus subunits of the heterotrimeric G protein that generate a high BRET signal when associated in their resting state. Upon receptor-induced G protein activation, the BRET signal decreased due to the dissociation of the α and βγ subunits. Mock cells (vehicle treated cells) were used as negative control and the mu opioid receptor (MOR) was used as a positive control in our assays, coupling to all Gα_i/o_ proteins as previously documented in the literature. We observed that WT ADGRV1β yielded a weak constitutive activation of the G_i_ protein, with quite similar coupling efficiencies to Gα_i1_, Gα_i2_ and Gα_i3_ (**Figure 1**). In contrast, no activation was detected for the G_i_-related Gα_oA_ or Gα_oB_ proteins, suggesting a selective coupling to G protein subtypes. The constitutive activity level of ADGRV1β is comparable to that of the μ-opioid receptor. Comparison of the results between MOR and ADGRV1 β receptors indicates that the selective coupling of ADGRV1β is not due to differences in G protein subunit expression. Interestingly, no coupling was observed for the G_q_ and G_s_ protein pathways, as measured by second messenger accumulation assays—cAMP HTRF® assays (*Revvity*) for G_s_ activation, or IP1-3 accumulation using IP-One HTRF® (REVVITY) for G_q_ activation (data not shown). This suggests that ADGRV1β couples exclusively to G_i_ proteins in our experimental conditions.

**Figure 1.**
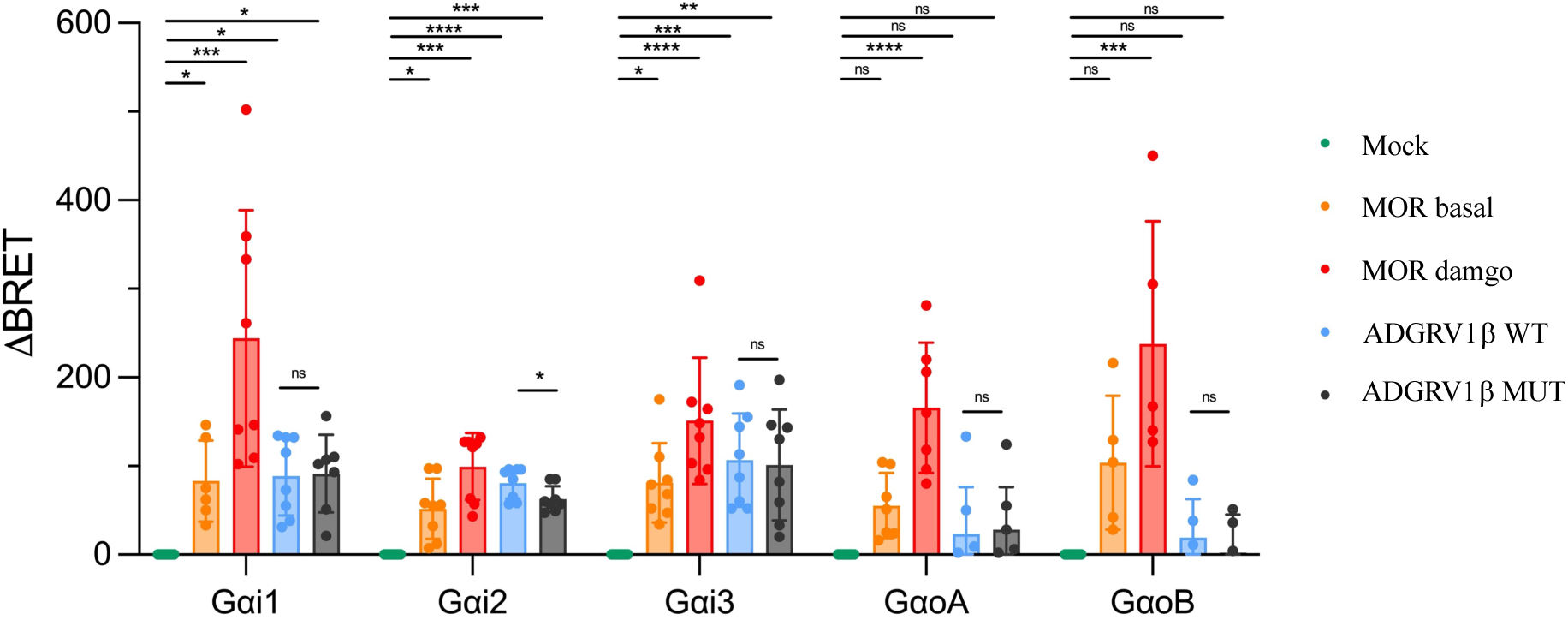
ADGRV1β constitutive signaling in cell. The constitutive signaling of ADGRV1β WT (TA-SVYAVYA) and MUT (TA-SVAAVAA) were assayed by BRET detection of the dissociation of Gα-RLuc and βγ-Venus subunits upon receptor-induced activation. Data are expressed as ΔBRET over Mock, where the Mock signal (vehicle treated cells) was subtracted to the agonist signal. The known Gαi/o-coupled receptor MOR was used as a positive control in absence and presence of its agonist damgo. Data are represented as mean +/-SD.

We then investigated whether the Gi coupling activity was mediated by the Stachel peptide. To this end, an ADGRV1 mutant was generated by substituting the residues at positions 3 and 6 (Y5886A and Y5889A, respectively) of the Stachel peptide, reported to be critical for its agonist function in other aGPCRs ^20^. Unexpectedly, this mutant ADGRV1β (MUT, of sequence -SVAAVAA-), harbouring both substitutions, exhibited Gi coupling activity comparable to that of the wild-type receptor. A partial reduction in Gα_i2_ coupling observed with ADGRV1β MUT suggests a minor agonist effect of the peptide. However, this variation is weak compared to the receptor’s inherent constitutive activity. This result indicates that the Stachel peptide is not responsible for mediating Gi coupling in ADGRV1. Conversely, the synthetic and selective agonist DAMGO ([D-Ala², NMe-Phe⁴, Gly-ol^5^]-enkephalin) activates the μ-opioid receptor regardless of its coupling to Gαi/o proteins. The expression at the cellular surface of the different proteins, MOR, ADGRV1β WT and MUT, has been previously controlled by ELISA assay in Human Embryonic Kidney 293 (HEK) cells with and without permeabilization (**Figure S2**). Therefore, we concluded that the observed signalling primarily results from the weak, yet significant, constitutive coupling of ADGRV1β. In order to further understand the molecular basis that may explain the lack of receptor activation by its agonist peptide, we conducted biophysical and preliminary cryo-Electron Microscopy (cryo-EM) studies of mouse ADGRV1β (Uniprot code B8JJE0), which was efficiently expressed in Sf9 cells and purified in its apo form in LMNG/CHS micelles to homogeneity.

### Biophysical characterization of ADGRV1β in LMNG/CHS micelles

Analytical ultracentrifugation (AUC) and small-angle X-ray scattering (SAXS) experiments were conducted on ADGRV1β to investigate its oligomeric state and shape in solution (**Figure 2**). We used the main elution peak of the gel filtration, the last step of purification, for biophysical characterization (**Figure 2A**). AUC data of ADGRV1β highlighted a main species at 4.0 S with a frictional ratio of 1.4 compatible with an elongated monomer (**Figure 2B, Table S1**) and a hydrodynamic radius of 5.3 nm. The dual UV/visible and interference detection system allowed the integration of the peaks at 280 nm and interference. The ratio of each protein and detergent in the peak could be resolved and revealed a mass of 111 kDa for the LMNG/CHS detergent surrounding the protein and an overall calculated mass of 158 kDa, compatible with the theoretical ADGRV1β molecular weight of 47.2 kDa. Consistent with the AUC results, the SAXS experiments (**Table S2**) showed a monodisperse distribution of monomers in solution (**Figure 2C-E**). The calculated molecular mass of 138 kDa derived from the extrapolated intensity I(0) at the origin is also consistent with the results of AUC experiments. The maximum distance of the protein and the radius of gyration values derived from the electron pair distance distribution function are 183 Å and 45 Å, respectively (**Figure 2D**). The Kratky plot (**Figure 2E**) showed a peak with a bell shape followed by an increase; this profile is characteristic of a compact domain with a flexible region and consistent with the structured 7TM domain embedded in the detergent micelle and the flexible C-terminal domain of ADGRV1β. Altogether, these results indicate that the protein is monomeric at the concentrations used for the rest of the study, with a globular domain embedded in the detergent micelle corresponding to the 7TM linked to the exposed, flexible region of the ICD.

**Figure 2.**
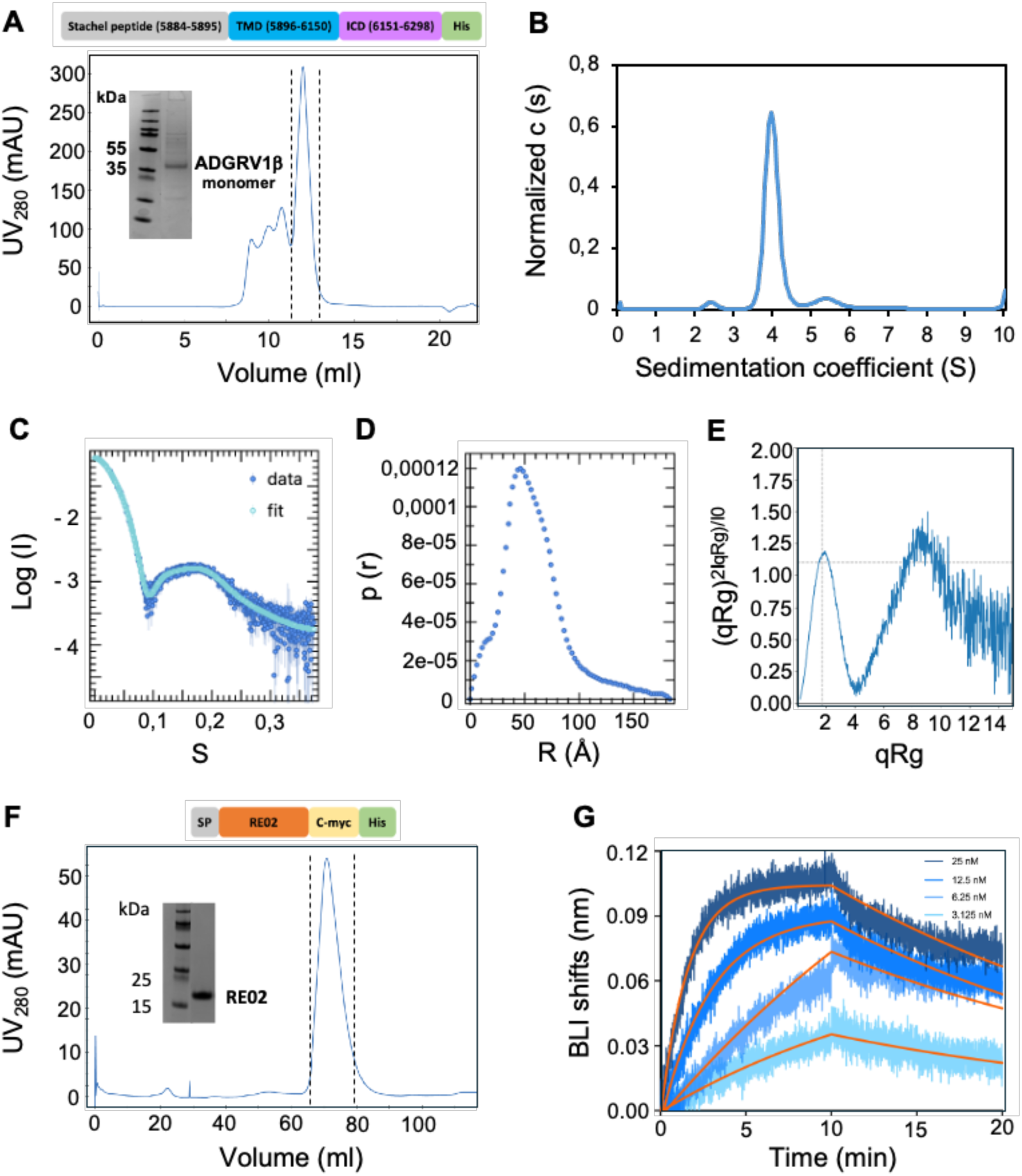
Biophysical properties of ADGRV1β and RE02 VHH. A) SEC profile and SDS-PAGE analysis of the purified ADGRV1β. B) Sedimentation coefficient distribution of ADGRV1β obtained by AUC. Data were obtained at 2 μM protein concentration. C) SAXS intensity of ADGRV1β in function of S. The fitted intensity is represented in cyan, computed with PRIMUS. D) Distribution of radius R (distance) of ADGRV1β with a maximal distance 183 Å. B. E) Dimensionless Kratky analysis of ADGRV1β. F) SEC profile and SDS-PAGE analysis of the purified RE02 VHH. G) BLI real-time monitoring of the interaction between immobilized ADGRV1β and varying concentrations of RE02 VHH in solution. The K_D_ value was obtained by the association/dissociation curves using a 1:1 binding model (orange lines).

### Nanobody engineering targeting ADGRV1β

Preliminary cryo-EM Single Particle Analysis (SPA) structural studies were carried out on ADGRV1β to solve its apo-state structure in LMNG/CHS detergent micelles. These experiments resulted in images of poor quality, essentially due to the dynamics of the transmembrane helices, which made it impossible to reconstruct the structure at high resolution. We therefore sought to identify nanobodies (VHHs) that target ADGRV1β, with the dual aim of reducing the intrinsic dynamics of its 7TM domain and providing additional extramembrane density to facilitate SPA.

Two alpacas were immunized with either purified ADGRV1β in LMNG/CHS or with HeLA cells transiently transfected with a cDNA coding for ADGRV1β, which is expressed at their surface. Indeed, injecting transfected cells may increase the chances of isolating conformation-specific VHHs in comparison with a detergent solubilized receptor ^27^. Pannings were performed on both librairies, and VHHs were selected that specifically bound to ADGRV1β, without cross-reactivity to the ICD domain. Notably, one VHH, designated RE02, was identified from both immunization approaches and proved to be a good ligand for ADGRV1 based on the titrations performed by ELISA. Therefore, RE02 was used for further experimental characterization in complex with ADGRV1β.

To optimize the formation of the VHH/ADGRV1β complex for structural characterization, we measured the affinity of RE02 for ADGRV1β. Large batches of pure RE02 were produced from *E. coli* extracts (**Figure 2F**). We used Bio Layer Interferometry (BLI) by coupling ADGRV1β to NTA Ni^2+^ biosensors and keeping the VHH in solution. RE02 displayed a dissociation constant (K_D_) value of 3 nM, indicating a very high affinity for the GPCR (**Figure 2G**). The calculated half-life time (T_1/2_) was estimated to be over 10 minutes, reflecting the high stability of the complex. Based on these properties, we decided to use RE02 to form a complex with ADGRV1β and determine its structure by cryo-EM.

### Cryo-EM structure of ADGRV1β bound to the RE02 nanobody

For GPCRs, low expression levels, conformational dynamics, and the frequent lack of available antagonists to stabilize them make structural determination of their inactive state particularly challenging. Therefore, protein fusions are routinely used like T4 lysozyme domain or other thermostable small proteins such as rigid BRIL (the thermostabilized apocytochrome b562a). These proteins are inserted into the third intracellular loops of GPCRs (ICL3) to increase their stability but also to ease structural determination by providing an extra-membrane density that aids particle alignment during SPA analysis. In designing our approach, we specifically aimed to avoid introducing any modifications into the GPCR sequence, to preserve the integrity of the intracellular loops of ADGRV1β. To achieve this, we used RE02 as a structural and stabilizing tool for the cryo-EM structure determination of ADGRV1β.

#### The cryo-EM structure of ADGRV1β/RE02 complex

For grid preparation, the complex was formed from both proteins purified separately and mixed with an optimized 10-fold molar excess of nanobody and a final concentration of ADGRV1 of 0.3 mg/ml. Using cryo-EM single-particle analysis, we solved the structure of ADGRV1β in complex with RE02 at a global resolution of 3.8 Å (**Figure 3A, Figure S3, Table S3**). The quality of the density map allowed us to unambiguously assign the 7TM core, the three extracellular loops (ECLs) and the three intracellular loops (ICLs) of ADGRV1β. Most side chains were built, except in a few regions where the density map was poorly resolved (**Table S4**), particularly in the lower halves of TMI and TMVII, which exhibit flexibility and dynamic behaviour. These regions are mainly exposed to the detergent micelle. Additionally, the extracellular N-terminus, including the Stachel peptide, and the intracellular cytoplasmic domain of ADGRV1β are not visible in our density map, likely due to their high flexibility.

**Figure 3.**
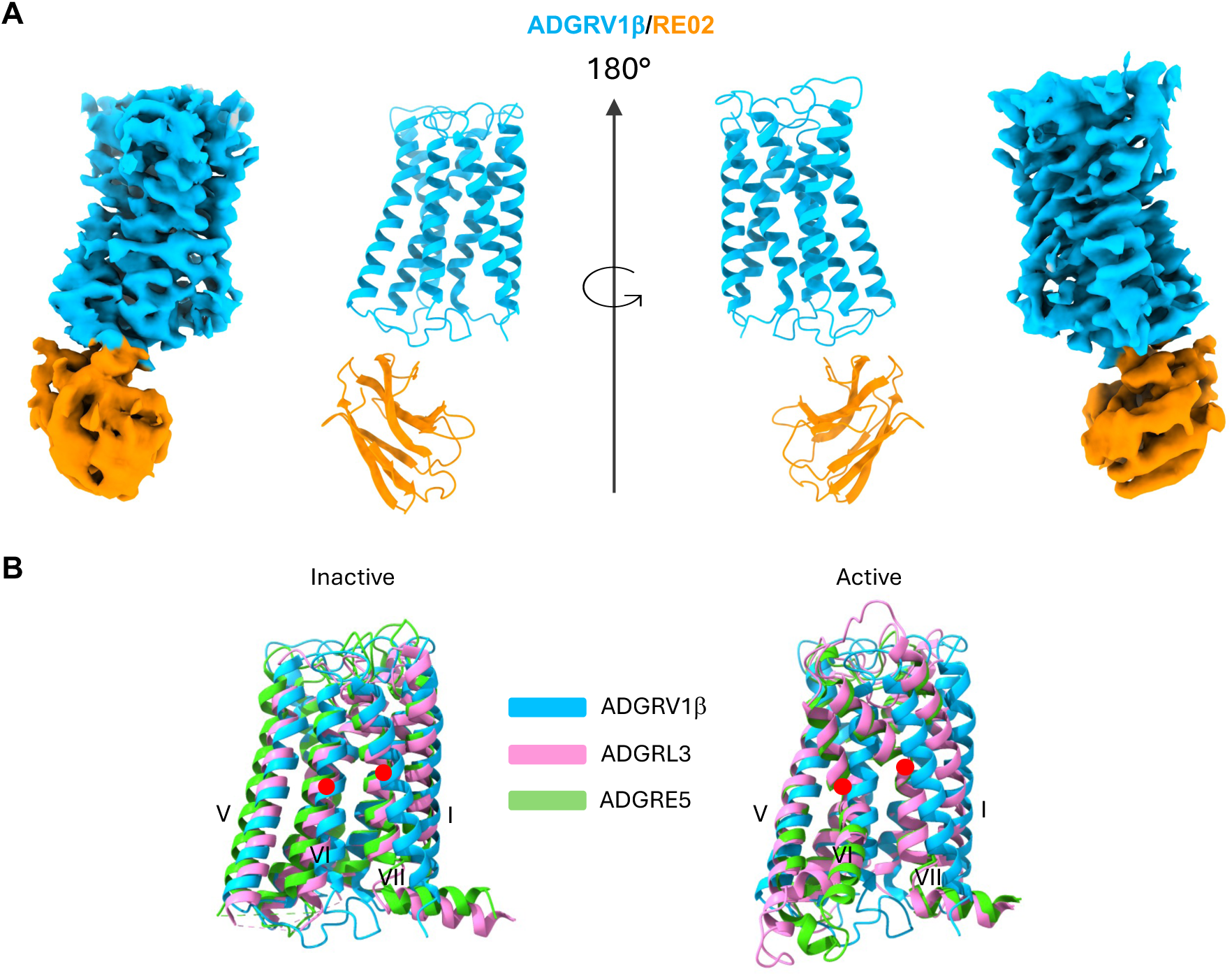
Structural state of ADGRV1β. A) Cryo-EM map and structure model of ADGRV1β (Blue) bound to RE02 VHH (orange). B) Superimposition of ADGRE5 (PDB: 8IKL) and ADGRL3 (PDB: 7SF7) structures with the structure of ADGRV1β in their inactive state (left) and in their active state (right, ADGRE5 (PDB:8IKJ) and ADGRL3 (PDB:8JMT)).

Regarding the VHH, most of the main chain of RE02, which is bound to the intracellular interface of ADGRV1β, was resolved. The RE02 region on the opposite side of the GPCR/VHH interface was poorly resolved (**Table S4**), primarily because a static mask focused on the 7TM was applied during the refinement process. This mask was necessary to resolve the GPCR region of ADGRV1β and the interface with RE02 at high resolution.

#### ADGRV1β binds to RE02 in its inactive state

Recently, the structures of ADGR-G1, -G2, -G3, -G4, -G5, -F1, -D1, -E5 and -L3 ^18 19 20 21 22 23 24 25^ in their active states have been determined by SPA in complexes with G or mini-G proteins, reporting the activation mechanism by the N-terminal Stachel peptide. The Stachel peptide of aGPCRs consists of 15-25 residues located upstream the TMI with the Tethered Agonist (TA) including approximately the first 7 residues (**Figure S4**). All structures share similar conformational insights with the TA adopting a short alpha helical conformation that binds to an orthosteric pocket unveiled by a groove formed between the tips of TMI, VI, VII on one side and TMII, III, V on the other. The most characteristic conformational signature of the active state of these aGPCRs lies in a kink within the helices TMVI and TMVII (**Figure S4**). This conformational transition creates a groove at the intracellular interface to accommodate the insertion of the Gα helix 5, thereby mediating the interaction with GPCRs.

To identify the conformational state of ADGRV1β bound to RE02, we compared our structure with ADGRE5 (CD97) and ADGRL3, the only two aGPCRs for which both active and inactive structures have been solved by SPA ^28 29^ (**Figure 3B**). The superimposition of our ADGRV1β structure with the active ADGRE5 (PDB 8IKL) and ADGRL3 (PDB 7SF7) structures reveals a clear absence of kinks in TMVI and TMVII in ADGRV1β (**Figure 3B, left**). The values of Cα Root Mean Square Deviation (RMSD) calculated for TMVI-TMVII are 3.2 Å and 2.4 Å, when compared to ADGRE5 and ADGRL3, respectively. Moreover, structural alignments of the inactive ADGRE5 (PDB 8IKJ) and ADGRL3 (PDB 8JMT) with ADGRV1β show a global similar conformation of the 7TM and an unambiguous straight conformation of their TMVI and TMVII (**Figure 3B, right**). However, TMVI and TMVII of ADGRV1 appear to be significantly shifted compared with those of the inactive ADGRE5 and ADGRL3 receptors (**Figure S5**). This shift may be explained by the adjustment of TMVI and TMVII conformations to accommodate the atypical, unusually long, and well-folded native ICL3 loop, which closely associates with the intracellular interface of ADGRV1 (**Figure S5**). In this comparison with inactive states, the Cα RMSD values for TMVI-TMVII are 1.9 Å and 0.9 Å for ADGRE5 and ADGRL3, respectively. Taken together, these structural insights and the absence of Stachel peptide density in the orthosteric pocket of our receptor confirm that we solved an inactive structure of ADGRV1β bound to RE02.

#### ADGRV1β misses part of the key elements for activation

Studies on other aGPCRs have characterized the Stachel peptide activation mechanism and revealed conserved regions critical for receptor activation: the Stachel peptide itself, its binding pocket, the ECL2 loop, the central hydrophobic core, and the helix kinks in TMVI and TMVII.

In the TA sequence of the Stachel peptide, three hydrophobic residues F^3^, L^6^ and M/L^7^ are highly conserved leading to the consensus motif (F^3^xxL^6^M/L^7^, superscripts indicate residue positions within the *Stachel* sequence) (**Figure 4A left**). These critical positions in helical conformation mediate similar interactions with the 7TM core in the reported *Stachel*-sequence-activated aGPCRs, highlighting their major importance in aGPCR activation (**Figure S4.B).** However, this consensus is not conserved in ADGRV1, which features a distinct TA sequence (Y^3^xxY^6^A^7^), preserved across 97 orthologues of the receptor (**Figure 4A right, S6**). Notably, the strong hydrophobic character of the three conserved residues in the TA sequence of aGPCRs is markedly reduced in ADGRV1.

**Figure 4.**
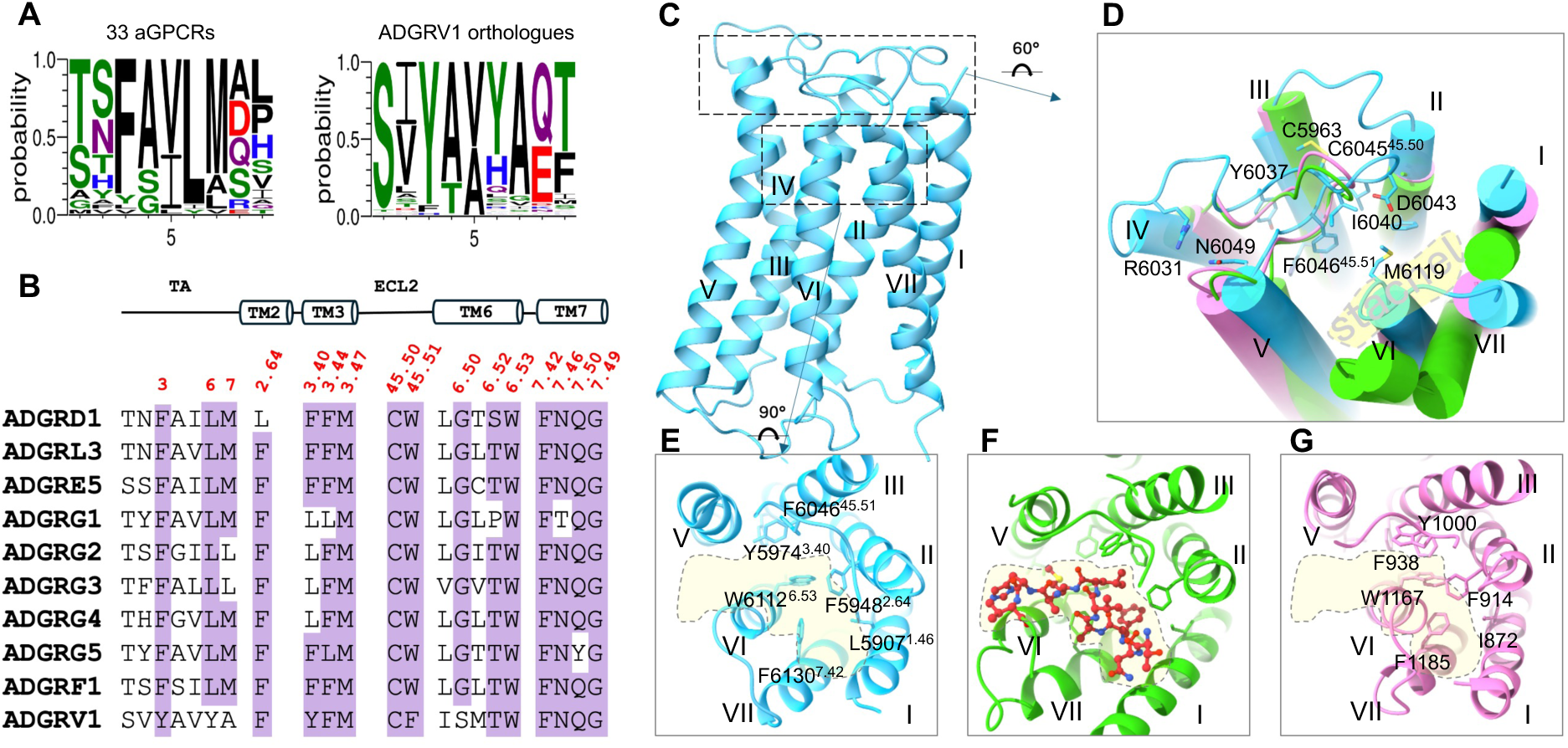
A) Logo representation of the nine first positions of the 33 aGPCR’s stachel peptide (left) and the 97 orthologues stachel peptide sequence of ADGRV1β (right). B) Sequence alignment of the TA stachel peptide and the most conserved binding pocket residues in aGPCR whose structure has been resolved. C) Global view of ADGRV1β structure (PDB: 9FTE, blue). D) Superimposition between ADGRV1β ECL2 (blue) and ADGRE5 ECL2 in its inactive (PDB: 8IKJ, pink) and active (PDB: 8IKL, green) states. E) Extracellular view of the ligand pocket of inactive ADGRV1β (PDB: 9FTE, blue) showing the critical residues of the inactive orthosteric site. F) Extracellular view of the ligand pocket of active ADGRL3 (PDB: 7SF7, green) with the stachel peptide TA represented in red. G) Extracellular view of the ligand pocket of inactive ADGRL3 (PDB: 8JMT, pink) showing the critical residues of the inactive orthosteric site.

In contrast, the residues in the orthosteric pocket that are crucial for the Stachel interaction and conserved in aGPCR are also present in ADGRV1(**Figure 4B, S7**). In active aGPCRs, the conserved hydrophobic residues of TA are inserted deeply in the bottom of the hydrophobic pocket, mainly interacting with the conserved residues F^2.64^, F/L^3.40^, W^45.51^, F^7.42^ and W^6.53^ (class B1 GPCR numbering in superscript ^30^). These critical residues of the 7TM interacting with the TA are also conserved in ADGRV1β. Indeed F5948^2.64^, Y5974^3.40^, F6046^45.51^, F6130^7.42^ and W6112^6.53^ contribute to a hydrophobic core including F5903^1.42^, L5907^1.46^ (**Figure 4C, E**). This hydrophobic pocket is highly conserved in the aGPCR inactive state illustrated by ADGRL3 in **Figure 4F, G**.

Another hydrophobic pocket in ADGRV1β is located downstream of W6112^6.53^, comprising F5978^3.44^, M5981^3.47^, L6065^5.46^, V6068^5.49^, F6105^6.46^, and I6108^6.49^. This cluster further strengthens interactions among TMIII, TMV, and TMVI. In other aGPCRs, three conserved residues—F^3.40^, F^3.44^, and M^3.47^ in TMIII form strong hydrophobic contacts with W^6.53^ in active aGPCRs. In ADGRV1, the corresponding residues Y5974^3.40^, F5978^3.44^, and M5981^3.47^ are present and conserved (Figure 4B, S7). In addition to these hydrophobic contacts, polar interactions also contribute to the stabilization of the active conformation. Two conserved polar residues, N^7.46^ and Q^7.49^ (forming the N^7.46^xxQ^7.49^ motif) (Figures 4B, S7), establish hydrogen bonds with the indole nitrogen of W^6.53^, a residue analogous to the “toggle switch” W^6.48^ in class A GPCRs ^31^.

In previously solved aGPCRs, the large extracellular loop 2 (ECL2) acts as a lid covering the orthosteric site. The ECL2 interacts primarily with the sixth TA residue, a conserved leucine (F^3^xx**L**^6^M/L^7^), through its conserved residue W^45.51^(Figure S4B). In ADGRV1β, the equivalent residue F6046^45.51^ also extends into the interior of the 7TM domain, contributing to the hydrophobic core of the orthosteric site in the inactive state (**Figure 4D**). The extended conformation of the ECL2 is also stabilized by hydrophobic interactions and a disulfide bond between the TMIII residue C5963^3.29^ and the ECL2 residue C6045^45.50^, two cysteine residues that are conserved within the aGPCR family (**Figure S7**). We then compared the conformation of the 21-residues ECL2 loop of ADGRV1β with that of the corresponding 16-residue loop in both the active and inactive states of ADGRE5 (**Figure 4D)**. We observed that the conformation of the ECL2 in ADGRE5 is similar in either its active or inactive forms and that the ECL2 of ADGRV1β and ADGRE5 align well. From these results, we conclude that the conformation of the ECL2 in ADGRV1β is likely not responsible for preventing the receptor’s activation by its tethered agonist peptide.

Finally, regarding the activation kinks, we observed a unique feature in ADGRV1. In other aGPCRs, the conserved W^6.53^ acts as a sensor of TA binding, initiating conformational kinks in TMVI and TMVII around G^6.50^ and G^7.50^, respectively (Figure S4). These kinks enable G protein binding at the intracellular face of active aGPCRs. The G^6.50^ and G^7.50^ residues, essential for helix unwinding in class B1 GPCRs, are also highly conserved in aGPCRs. Mutations at these key positions in several aGPCRs have been shown to significantly impair receptor activity ^18–25^. In ADGRV1, while G6140^7.50^ is conserved in ADGRV1, the glycine at position 6.50 is unexpectedly replaced by a serine (S6109), which may influence the degree of kink formation in TM6 (**Figure 4B**).

Altogether, we showed that ADGRV1 shares most critical residues and motif elements necessary for the activation mechanism shown for other aGPCRs but lacks the consensus Stachel peptide sequence and G^6.50^ that are key actors of aGPCRs activation. As we observed a weak constitutive activity of ADGRV1 in cellular assays, we hypothesize that those two misconserved elements could impair its activation.

### The ADGRV1β ICL3 conformation and the binding interface with the nanobody RE02

In the inactive structures of ADGRE5 and ADGRL3, the native conformation of ICL3 is not described because of a BRIL domain insertion within their ICL3 loops. In our structure, we successfully reconstructed the large intracellular loop 3 (ICL3, 17 residues) of ADGRV1β, which is the first reported native ICL3 conformation in the inactive state of an aGPCR. In ADGRV1β, ICL3 acts as a lid predominantly covering the entire intracellular interface of the protein (**Figure 5A, B**). Analysis of this region reveals a hydrophobic core involving W6082, V6088, and F6089 from the ICL3. Their side chains point toward the transmembrane domain, contacting hydrophobic residues from TMII to TMVII, F5929, F5988, F6072 and L6148, that surround the ICL3 (**Figure 5C**). Therefore, the wide opening in active structures to accommodate the G protein is closed and buried by the ICL3 in our ADGRV1β inactive state.

**Figure 5.**
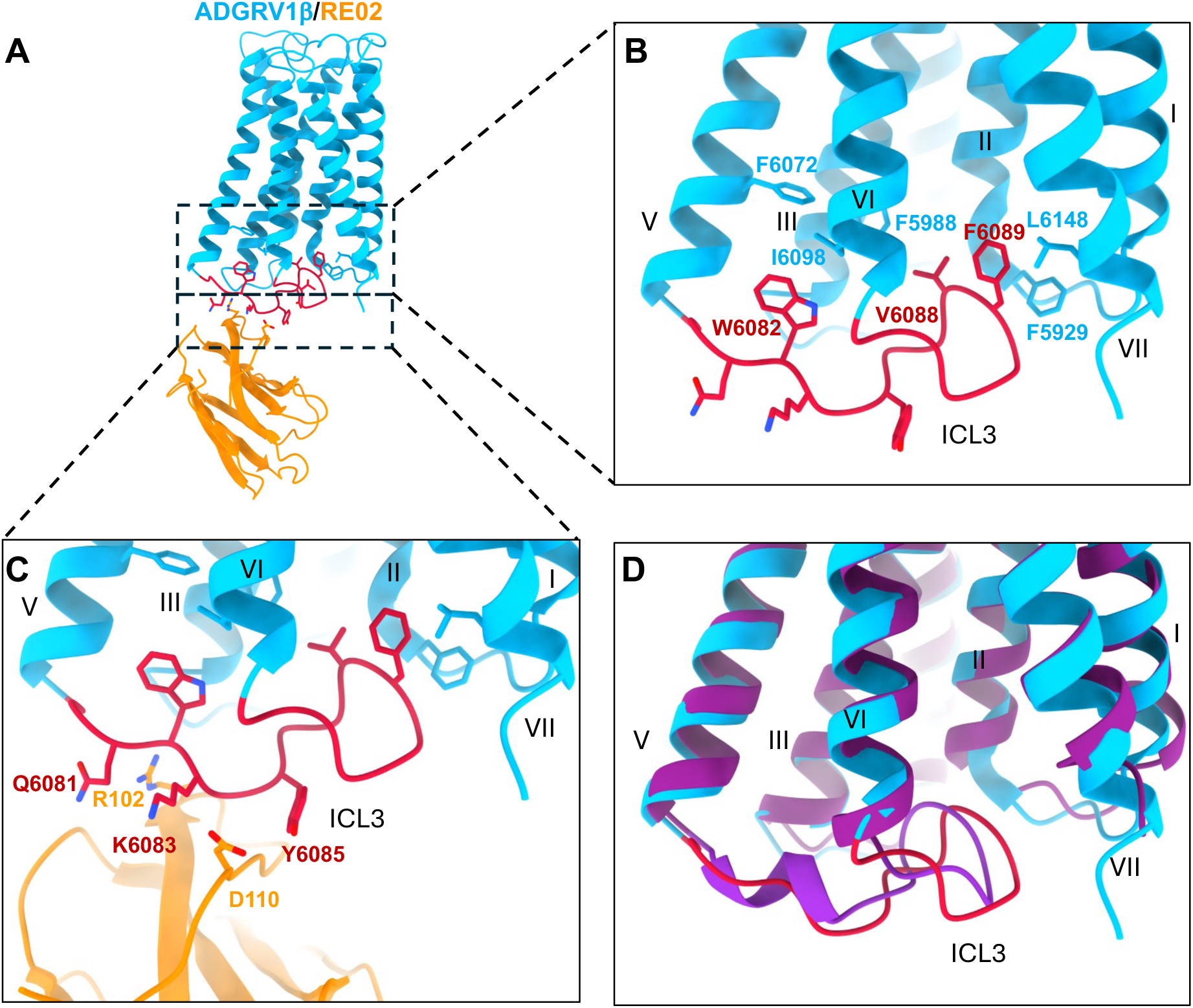
A) Structure representation of ADGRV1β (blue) bound to RE02 VHH (orange). B) View of ICL3 conformation within the TMD of ADGRV1β. C) View of the ADGRV1β /RE02 interface. D) Superimposition of ADGRV1β EM structure with its Alphafold 3 prediction showing ICL3 conformation.

The RE02 VHH binds to ADGRV1β through its CDR3, which contacts the receptor’s ICL3 (**Figure 5A, C**). The complex is mainly stabilized by several polar interactions. The D110 residue of RE02 is engaged in polar interactions with Y6085, and forms a salt bridge with K6083, from the ICL3. Additionally, a polar interaction is observed between R102 and Q6081 (**Figure 5C**). Collectively, these interactions significantly contribute to the binding affinity in the nanomolar range of RE02 for ADGRV1β. It is worth noting that the atypical conformation of the ICL3 we modelled from our density map is very similar to the one predicted by AlphaFold3 (**Figure 5D**).

### Molecular Dynamics simulations of ADGRV1β

To assess the effect of the VHH on the structural and dynamic properties of ADGRV1β, we conducted atomistic Molecular Dynamics (MD) simulations of ADGRV1β starting from our SPA structure, both in presence and absence of the VHH. All simulations were performed in explicit water and POPC lipid bilayer. For each construct, we conducted three replicate simulations, each lasting 2 µs, resulting in a total aggregated simulation time of 12 µs (**Table S5**). We first verified that the global conformation of ADGRV1β determined by cryo-EM remained stable over the microsecond timescale in the absence of VHH. Therefore, we calculated the backbone RMSD from the cryo-EM structure of ADGRV1β (excluding the VHH and ICL3) during the course of the MD simulation (**Figure 6A**). In all three replicates, the ADGRV1 structure without VHH (**Figure 6A**, shades of red) exhibited variations over time relative to the initial structure that are similar to those observed in the ADGRV1-VHH complex (**Figure 6A**, shades of green). This behaviour indicates that the ADGRV1 structure solved by cryo-EM remains stable over the microsecond timescale in the absence of VHH, and then, suggests that the nanobody does not have a significant effect on the structural and dynamic properties of ADGRV1β. We also observed that in all three replicate simulations of the ADGRV1/VHH complex, the nanobody was extremely dynamic, with one instance where the VHH detached from ADGRV1 after a few nanoseconds of simulation.

**Figure 6.**
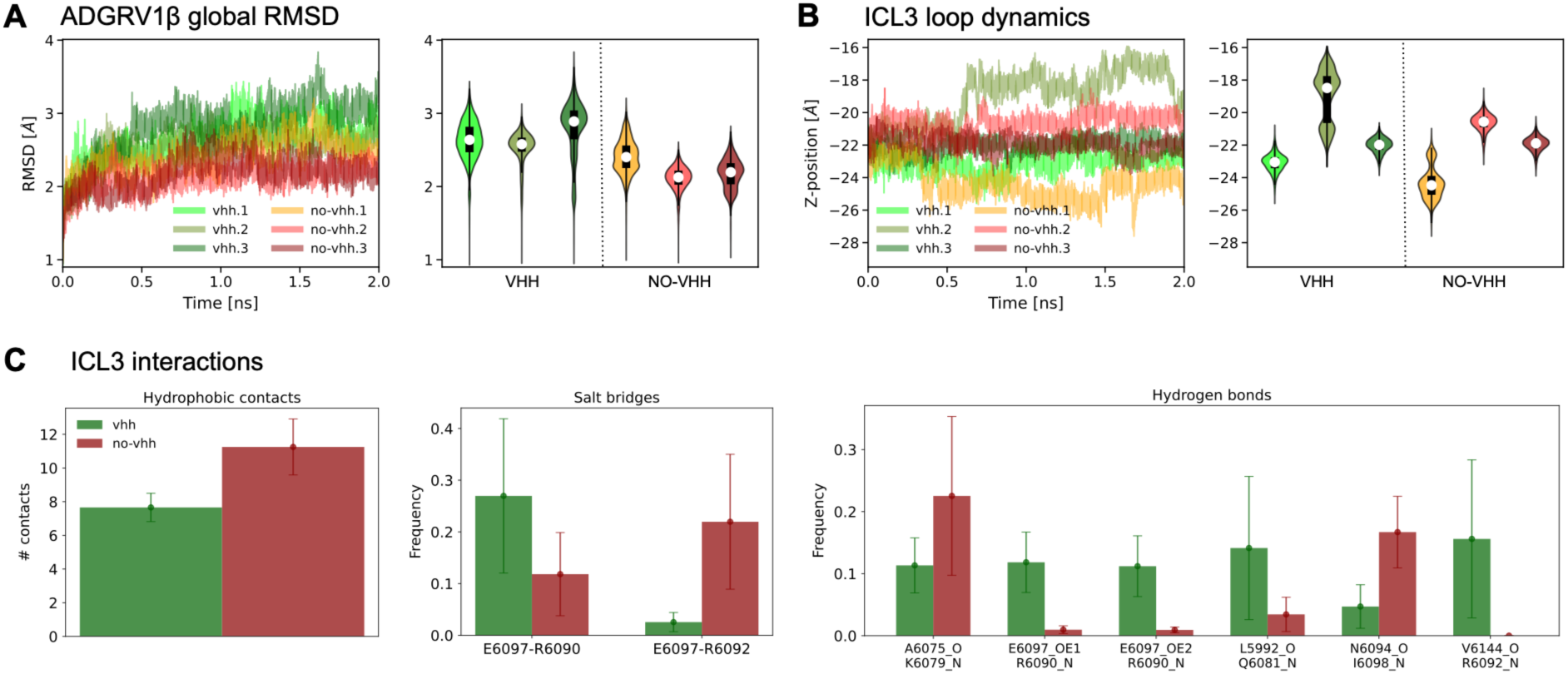
Analysis of the MD simulations. A) Backbone r.m.s.d. from the cryo-EM structure of ADGRV1β during the course of the MD simulation in presence (green shades) and absence (red shades) of the VHH (left panel). r.m.s.d. violin plots for each simulation (right panel). B) Z-coordinate of the ICL3 loop backbone atoms during the course of the MD simulation in presence (green shades) and absence (red shades) of the VHH (left panel). Z-coordinate violin plots for each simulation (right panel). In the violin plots of panels (A) and (B), the white circle corresponds to the median value, the black rectangle extends from the first to the third quantiles, and the thin black line represents the 95% confidence intervals. C) Analysis of the interaction between the ILC3 loop and the rest of ADGRV1β in presence (green) and absence (red) of the VHH: mean number of hydrophobic contacts formed during the course of the simulation (left panel), and frequency of formation of salt bridges (centre panel) and hydrogen bonds (right panel). The bar plots represent the average value across the three replicate simulations, and the error bars the variation across replicates.

We then focused our analysis on the ICL3. Since the VHH binds to ADGRV1 in the ICL3 region, we wondered if the nanobody could influence the position, structure, and dynamic of this loop. Specifically, we investigated whether the buried position of the ICL3 within the TM region of ADGRV1β was due to the presence of the VHH, which could be blocking access to the bulk solvent. Our analysis indicates that ICL3 remains buried within the TM region even in the absence of the VHH (**Figure 6B**). These conformations are stabilized by hydrophobic contacts, as well as weak, transient hydrogen bonds and salt bridges between the ICL3 and the TM region of ADGRV1β (**Figure 6C**). We therefore concluded that the buried conformation of the ICL3 observed by cryo-EM in presence of the nanobody might remain stable even in its absence. This suggests that the ICL3 could potentially compete with G-protein binding, regulating ADGRV1β activation.

In addition, Alphafold3 structural predictions of ADGRV1β systematically result in an inactive form of ADGRV1β, which is consistent with our SPA structure. The two structures are very similar with a RMSD of 1.5 Å. Interestingly, in the AlphaFold3 model, the ICL3 exhibits a comparable lid conformation with a local RMSD of 2.6 Å to the SPA structure, largely covering the intracellular surface of the 7TM (**Figure 5C**). These observations are consistent with the results of our MD simulations, which show that the conformation of the ICL3 is packed against the 7TM in the absence of the VHH.

## Discussion

Adhesion G protein-coupled receptors (aGPCRs, class B2) represent a fascinating class of GPCRs that play critical roles in regulating cellular interactions, including adhesion, polarity, and cell migration. Despite their physiological and pathological significance in various immune, cardiovascular, neurological processes, the activation mechanisms of these receptors remained elusive for a long time due to their structural complexity. This complexity is characterized by the presence of numerous domains which compose their large extracellular region, and their unique autoproteolytic ability which cleaves these GPCRs in two subunits. Over the past three years, remarkable progress has been made in elucidating their structures, primarily due to major advances in cryo-EM. These studies have revealed unique activation mechanisms for nine receptors spanning aGPCR subgroups, notably highlighting the role of the Stachel peptide, which acts as an intramolecular agonist to trigger aGPCR signalling. Most of these aGPCRs structures have been solved in their active form in complex with their specific G protein. While the conformational transitions of GPCR classes A, B1, C, and F have been well characterized ^32^, the recent characterization of the first inactive structures of aGPCRs, specifically ADGRE5 and ADGRL3, by cryo-EM has provided a more comprehensive understanding of the aGPCR activation process. The lack of inactive structures certainly highlights the difficulty of studying these rather small and dynamic GPCR domain.

ADGRV1 is the only receptor representing the IX sub-family of aGPCRs, with no structural data available to date. Although its signalling activity has been studied in various cellular models showing the preferential coupling of its CTF domain to G_i_ protein, the activation mechanism is still controversial, particularly regarding the role of its Stachel peptide. In this article, we present the first high-resolution structure of ADGRV1-CTF (ADGRV1β) in its inactive state, in complex with RE02, a nanobody (VHH) specific to the receptor. We combined our structural data with the functional analysis of ADGRV1β activity in cell. Our inactive structure of ADGRV1 is the first to provide insights into the CTF domain of an aGPCR without the use of antagonistic molecules or sequence modifications. This approach enabled us to capture the native inactive structure of an aGPCR, detailing its seven 7TM as well as all extracellular and intracellular loops, particularly the ICL3. Although the extracellular region encompassing the TA and the entire C-terminal cytoplasmic domain of ADGRV1β are not seen in the density map, all the loops and helices are well defined in our structure.

When compared with other inactive aGPCR structures, ADGRV1 similarly exhibits a compact orthosteric pocket in the absence of Stachel peptide binding, stabilized by a network of hydrophobic interactions. This inactive pocket in ADGRV1 is primarily formed by highly conserved hydrophobic residues, including F5948^2.64^, F6046^45.51^, Y5974^3.40^, F5978^3.44^, F6130^7.42^, and W6112^6.53^. This tryptophan is the structural equivalent of the conserved ‘toggle switch’ W6.48 in class A GPCRs, which plays a crucial role in the conformational transition of adhesion GPCRs toward their active state. The adhesion GPCRs whose structures have been solved are typically activated by their tethered agonist peptide adopting a helical conformation in the orthosteric pocket. TA is defined by the consensus motif F^3^xxL^6^M/L^7^. In such cases, W6.53 acts as a sensor for the binding of this sequence. Around this critical residue, significant conformational changes occur, particularly involving the conserved residues F^3.40^, F^3.44^, M^3.47^, and the N^7.46^xxQ^7.49^ motif.

While the critical positions for Stachel peptide interaction are conserved within the orthosteric pocket of ADGRV1, the receptor itself features a non-conserved TA sequence, with a substitution of key residues involved in activation (Y^3^xxY^6^A^7^). These substitutions result in the peptide having a globally less hydrophobic character compared to the consensus sequence (F^3^xxF^6^M^7^). Thus, we expect a weaker interaction between the conserved orthosteric pocket of ADGRV1 and the non-consensus sequence of its Stachel peptide compared to what is observed for others aGPCRs. ADGRV1 has the Stachel peptide sequence that is the most divergent from the accepted consensus amongst all aGPCRs. The highly conserved G^6.50^ and G^7.50^ residues, also found in class B1 GPCRs ^33^, are essential for the conformational transition inducing kinks in the middle of TMVI and TMVII, which lead to G protein engagement and activation. Furthermore, ADGRV1 is the only aGPCR that lacks a glycine (G) at position 6.50. Such a substitution by a serine (S6109) could reduce the local helical flexibility, thereby preventing the transition toward an active conformation. Together, these features of ADGRV1β may explain our findings in cells, which demonstrate its weak constitutive activity and the inability of the Stachel peptide to enhance this activity.

Our results are consistent with the recent work of Dates et al.(2024)^17^ who assessed the TA-depending activities of all 33 aGPCRs through cellular assays. The authors also conclude to the heterogeneity in the activation mechanisms of aGPCRs, showing that only 50% of these receptors are activated by their Stachel peptide. Most receptors that are not activated by their Stachel peptide, including ADGRV1, lack the G6.50 residue on their TMVI, indicating a potential link between this position and the Stachel-mediated activation mechanism. Nevertheless, our results also confirm a preferential coupling of ADGRV1 to the G_i_ protein, which is consistent with previous published results ^15^. Taken together, our structural and functional results challenge the role of the TA-Stachel peptide in ADGRV1 activation and suggest that its mechanism is largely distinct from the activation model proposed for other aGPCRs studied to date.

Another unique features of ADGRV1β concerns the sequence and the conformation of its ICL3 that tethers TMV to TMVI. This long sequence adopts a lid-like conformation, covering and obstructing the intracellular interface of ADGRV1 through a series of hydrophobic interactions with the 7TM domain. The stability of this ICL3 conformation, assessed through our MD simulations, represents another feature of ADGRV1 that may affect ADGRV1β activation by competing with G-protein binding. Previous studies on GPCR β_2_AR support the notion that a long or structured ICL3 can function as an intrinsic negative regulator of GPCRs, and can tune signalling specificity by inhibiting receptor binding to G protein subtypes that weakly couple to the receptor ^34^.

The activation mechanisms of aGPCRs have only recently begun to be investigated. To date, nine aGPCRs have been studied, all of which appear to share a similar molecular activation mechanism that depends on the Stachel peptide, regardless of the specific G protein involved. In contrast, our study shows that ADGRV1β activation is independent of its Stachel peptide. We attribute this non-canonical behaviour to several unique structural and sequence features of the receptor. The extracellular domains of ADGRV1 appear to play a role in the receptor activity, regulating the G protein coupling profile to the CTF ^4 5 6^, but the molecular mechanisms remain unknown. Understanding the molecular mechanisms of receptor activation is crucial for grasping better insight into its role in cellular signalling within sensory cells and its involvement in related pathologies. Several potential pathological missense mutations have been identified in the CTF of ADGRV1β, establishing a potential link between the receptor activity, its function in hair cells, and possible deafness pathologies. The search for agonists will be essential to understand the activation of ADGRV1, as well as to investigate other possible downstream signalling pathways, such as non-canonical pathways in sensory cells.

## Materials and Methods

### Reagents and plasmids for in cellulo signalling studies

ADGRV1β wild-type or mutated ADGRV1β nucleotidic sequence synthesis and insertion into the pRK5 vector were ordered from Genecust (France). In the mutated ADGRV1β peptide sequence, the TA Stachel peptide SV<u>Y</u>AV<u>Y</u>A was replaced by SV<u>A</u>AV<u>A</u>A. Both constructs were designed to contain a Flag-tag at the intracellular C-terminal end. Coelenterazine-h (S2011) was purchased from Promega Corporation (Madison, WI, USA).

### Cell culture and transfection

HEK293 cells were obtained from ATCC (CRL-1573, Manassas, VA, USA). HEK293T cells were cultured in DMEM (Thermo Fisher Scientific) supplemented with 10% foetal bovine serum (Sigma-Aldrich, Saint-Quentin-Fallavier, France) and incubated at 37°C in a CO_2_ incubator. Unless stated differently, HEK293 cells were transfected using Lipofectamine® LTX with Plus™ Reagent (Thermo Fisher Scientific) in 96-well F-bottom black or white plates (Greiner Bio-One). 50000 cells were seeded with 100 µL DMEM in each well 24h before transfection in order to get cells at 70% confluence the next day.

### ELISA assay for receptor expression measurements

ELISA experiments were performed as previously described ^35^. Briefly, cells were fixed with 4% paraformaldehyde and blocked with phosphate-buffered saline (PBS) containing 1% foetal calf serum and then incubated 30 min at 0.5 mg/L with monoclonal anti-Flag M2 antibodies (SIGMA, L’isle-D’Abeau, Saint-Quentin Fallavier, France), anti-HA antibodies (clone 3F10, Roche Applied Science, Basel, Switzerland). When these primary antibodies were not conjugated with horseradish peroxidase (HRP), cells were further washed and incubated (30 min) with HRP-conjugated goat anti-rat IgG (0.5 mg/L, Jackson ImmunoResearch laboratories, Westgrove, PA, ISA) or anti-rabbit IgG or anti-mouse IgG (0.5 mg/l, Amersham Biosciences GE Healthcare, Chicago, IL, USA) for 30 min. After washes, bound antibody was detected by chemiluminescence using SuperSignal substrate (Pierce, Rockford, IL, USA) and a Mithras LB 940 plate reader (Berthold Biotechnologies, Bad Wildbad, Germany). As a control for intracellular ELISA quantification, cells were permeabilized for 5 min with 0.01% Triton X-100 just after being fixed.

### G protein BRET assay

Cells were plated at density of 50000 cells/well and reverse transfected: with the indicated ADGRV1β and the α-RLuc8 and βγ-Venus G protein subunits. The next day, cells were washed with PBS twice and incubated 10 min at 37 °C. RLuc signal intensity was measured in Mithras LB 940 Multimode Microplate Reader (Berthold Technologies) by measuring fluorescence emitted at 535 nm. Then, 5 µM coelenterazine was added and BRET measurements were done during 500 s, before the indicated compounds were added. BRET signal was then measured for 1500 s. Data was expressed as ΔBRET over basal, where the basal signal (vehicle treated cells) was subtracted to the agonist signal. miliBRET was calculated as the signal obtained at 535 nm over the signal obtained at 485 nm and adjusted by subtracting the ratios obtained when RLuc fusion proteins were expressed alone, and by multiply the result by 1000.

### Statistical analysis

All data analysis was performed using Prism 9 (GraphPad Software Inc., San Diego, CA, USA). Data are expressed as mean ± SD. The experiments were performed in triplicate and are representative of 5 to 8 independently performed experiments: one-way ANOVA or Brown-Forsythe and Welch followed by the post hoc Dunnett multiple comparison. p< 0.05 has been defined as a significant difference for all statistical analysis. Statistical analyses were performed with GraphPad Prism 9 (GraphPad software Inc, San Diego, CA, USA).

### ADGRV1β expression and purification

The cDNA sequence encoding murine long isoform ADGRV1β (aa 5883-6298) (Uniprot: B8JJE0_MOUSE) was chemically synthesized (Genscript). For protein purification, the sequence was modified to include an 8x Histidine tag at the very C-terminus of the receptor. The sequence was cloned in pFastBac1 plasmid for insertion by recombination (DH10Bac E. coli cells) into baculovirus shuttle vector (Bacmid) using the Bac-to-Bac Baculovirus Expression System (Thermofisher). High-titre recombinant baculovirus was produced according to the same manual. To express the protein, Sf9 insect cells were used and cultured with ESF921 medium (Expression Systems). The high-titre recombinant baculovirus P2 generated was used to infect insect cells grown to a density of 3×10 ^6^ cells per ml with an MOI of 2.5. The cells were collected by centrifugation after 48h of infection, and the cell pellets were stored at -80°C.

The pellets expressing ADGRV1β were resuspended in 50mM Tris HCl pH 7.5, 300mM NaCl, 0.2mM TCEP supplemented with Complete™, EDTA-free Protease Inhibitor Cocktail (Roche) then lysed by an Ultra-turrax. The membrane fraction was collected following centrifugation at 41000 rpm for 2h. The protein was then solubilized with 1% LMNG/0,1% CHS (Anatrace) for 1h at 4°C after Dounce homogenization. The solubilized fraction was collected following a centrifugation at 41000 rpm for 30 min and then loaded on an Ni++ metal affinity chromatography Hitrap Fast Flow column 1 ml (Cytiva) equilibrated in 50mM Tris HCl pH 7.5, 300mM NaCl, 0.2mM TCEP, 0,1% LMNG/0,01 CHS and 20mM imidazole. The column was then washed with 30 column volumes of 50mM Tris HCl pH 7.5, 300mM NaCl, 0.2mM TCEP, 0,01% LMNG 0,001 CHS and 40mM imidazole. ADGRV1β was eluted with 10 column volumes of 50mM Tris HCl pH 7.5, 300mM NaCl, 0.2mM TCEP, 0,01% LMNG/0,001 CHS and 250mM imidazole and then purified by size exclusion chromatography (SEC) on a Superdex 200 10/300 increase column (Cytiva) pre-equilibrated with 50mM Tris HCl pH 7.5, 150mM NaCl, 0.2mM TCEP, 0,00075% LMNG/0,000075 CHS. The fractions corresponding to the protein elution were pooled and concentrated to 1,5mg/ml (for EM experiments).

### ADGRV1β cytoplasmic domain expression and purification

The murine long isoform ADGRV1β cytoplasmic domain (ADGRV1β-cyto) (CCDS36737; 6149 – 6298) was expressed in fusion with a N-terminal Histidine tag followed by a TEV protease site. Protein production was performed in E. coli BL21 DE3 in LB, induced with 1 mM IPTG for 3 h30min at 20 °C. Lysis was done in Tris HCl pH 7.5 50 mM, NaCl 300 mM, 0.5 mM TCEP, supplemented with Complete™, EDTA-free Protease Inhibitor Cocktail (Roche). The cell lysis was performed on a Cell disruptor at 1.3 kbar. After 1 h centrifugation at 20000g, the supernatant was loaded on an Ni^2+^ metal affinity chromatography Hitrap Fast Flow column 5 ml (Cytiva) equilibrated in 50mM Tris HCl pH 7.5, 150mM NaCl, 0.2mM TCEP and 20mM imidazole. The column was then washed with 30 column volumes of 50mM Tris HCl pH 7.5, 150mM NaCl, 0.2mM TCEP, and 40mM imidazole. ADGRV1β-cyto was eluted with 10 column volumes of 50mM Tris HCl pH 7.5, 150mM NaCl, 0.2mM TCEP, and 250mM imidazole. The protein was subsequently eluted, and a last step of purification was performed by SEC using a Sephacryl S-100 HP 16/60 column (Cytiva).

### Analytical Ultracentrifugation (AUC)

Sedimentation velocity experiments on ADGRV1β were performed using an analytical ultracentrifuge (Beckman Coulter Optima AUC) equipped with an eight-hole AN60-Ti rotor with 12 mm six sectors centrepieces. The temperature was fixed at 4°C. The protein sample was prepared at 2 μM in 50mM Tris HCl pH 7.5, 300mM NaCl, 0.001% LMNG/0.0001% CHS, 0.5 mM TCEP and then centrifuged for 18h at 42000 rpm. Sedimentaion profiles were recorded by absorbance at 280nm and interference. Data were analysed with SEDFIT 15.1 using a continuous size distribution c(S) model^36^. The amount of detergent and protein in the main peak was determined by using the absorbance and interference signal as proposed by others ^37^. Values used for the calculations are presented in Supplementary Table S1.

### Small Angle X-ray Scattering (SAXS)

Synchrotron radiation X-ray scattering data were collected at the SWING beamline at Synchrotron Soleil (France) using the online HPLC system. 50 µl of purified ADGRV1β were injected at a concentration of 60 μM into a size exclusion column (Superdex 200 increase 5/ 150 Cytiva) in line with the SAXS flow-through capillary cell at 0.2 ml/min flow rate. The experiment was performed in a buffer containing 50mM Tris HCl pH 7.5, 300mM NaCl, 0,001% LMNG/0,0001% CHS, 0.5mM TCEP.

Initial data processing was performed using RAW and PRIMUS. The radius of gyration was evaluated using the Guinier approximation. The Dmax was determined from the distance distribution function P(r) obtained with the program GNOM. Additional details of the SAXS data collection are reported in Supplementary **Table S2.**

### Alpaca immunization

Animal experimentation was executed according to the French legislation and in compliance with the European Communities Council Directives (2010/63/UE, French Law 2013-118, February 6, 2013). The Animal Experimentation Ethics Committee of Pasteur Institute (CETEA 89) approved this study (2020-27412). Two young adult male alpacas (*Lama pacos*) were immunized on days 0, 21, and 28 with approximately 10^8^ transiently transfected HEK293 cells expressing ADGRV1β for the first alpaca, and the with purified protein for the second one. The immunogen was mixed with Freund’s complete adjuvant for the initial immunization and with Freund’s incomplete adjuvant for subsequent immunizations. The immune response was monitored by titrating serum samples by ELISA. Microplates were coated with either ADGRV1β purified protein, transiently transfected HEK cells, or the isolated intracellular domain of ADGRV1β. This allowed for detection of antibodies binding specifically to the GPCR domain of the protein. The antibodies were detected with polyclonal rabbit anti-alpaca IgGs. A blood sample of approximately 300 mL was collected from each immunized animal, and a Phage-Display library was constructed in a pHEN6 phagemid vector, comprising approximately 2 × 10^8^ different clones, as described before ^27^.

### RE02 nanobody expression and purification

The pHEN6 vector used for VHH library construction in *E. coli* TG1 strain, enables the expression of VHHs as recombinant proteins in *E. coli* BL21 (DE3) strains. This vector encodes a VHH fused to a C-terminal cMyc tag and a His tag, facilitating both detection and purification of the protein. Following overnight induction in 2YT broth with 1 mM IPTG at 30 °C, bacteria were harvested by centrifugation. The resulting pellet was resuspended in 1X PBS buffer supplemented with Complete™, EDTA-free Protease Inhibitor Cocktail (Roche), and periplasmic extraction was performed either by addition of 1mg/ml of polymyxin B sulfate (Sigma-Aldrich, # 5291-1GM) and incubation at 4 °C for 1 h with gentle shaking, or by osmotic shock in the presence of 200mM of sucrose. Periplasmic extracts were obtained by centrifugation at 12000 rpm for 45 min. The VHH protein was purified using a Co++ metal affinity chromatography HiTrap TALON^®^ crude 1 mL column (Cytiva). The column was washed with 40x column volumes of 1X PBS containing 10mM imidazole, then the protein was eluted in 1X PBS containing 500mM imidazole, followed by a Size Exclusion Chromatography (SEC) using a HiPrep^™^ 16/60 Sephacryl^®^ S-100 HR column pre-equilibrated with 1X PBS. The fractions containing the VHH were pooled and concentrated to 1 mg/ml for Electron Microscopy (Em) experiments. For BLI experiments, a TEV-cleavage of the RE02 his-tag was performed in 1X PBS.

### Real-time characterization of the RE02/ADGRV1β interaction by BLI

Biolayer interferometry assays were performed using NTA biosensors (18-5101, Sartorius) on an Octet Red384 instrument (Sartorius) at 25°C. All assays were performed using standard half-well black 96-well microtiter plates (655209, Greiner) with a volume of 120 μl per well. PBS buffer containing 1 mg/ml BSA (PBS-BT) was used for sensor pre-equilibration, baseline measurements, dissociation steps, and diluting the RE02 VHH. ADGRV1β was captured on the sensors via its His tag at 20 μg/ml for 1800 s. RE02 association was monitored for 1200 s in PBS-BT with concentrations ranging from 25 nM to 3,125 nM followed by dissociation for 1200 s in PBS-BT. Two wells containing only buffer instead of RE02 were assigned as reference wells. Signals from reference wells and reference biosensors devoid of ADGRV1β were subtracted, and curve fitting was performed using a global 1:1 binding model in the HT Data analysis software 11.1 (Sartorius, allowing to determine K_D_ values.

### Cryo-EM sample preparation

The ADGRV1β/RE02 complex was prepared immediately before grid preparation by mixing the two proteins, purified separately, to final concentrations of 6 μM for ADGRV1β and a 10-fold molar excess of nanobody. UltrAuFoil 300 mesh 1.2/1.3 grids were glow-discharged in a PELCO easiGlow at 25 mA for 15s. Subsequently, 3 µL of the sample was applied to the grids, blotted (Whatman filter paper, Grade 595) for 4s at 100% humidity and 12°C, and vitrified by plunge-freezing in liquid ethane cooled by liquid nitrogen using a Vitrobot Mark IV (Thermo Fisher). The vitrified grids were stored in liquid nitrogen until use.

### Cryo-EM data acquisition and processing

For grid screening, we used a Glacios microscope operated at 200kV equipped with a Falcon 4i detector (Thermo Fisher Scientific) at the Nanoimaging Core Facility of Institut Pasteur (Paris, France). Full cryo-EM datasets were collected with EPU version (ThermoFisher) on a Titan Krios G3 microscope operated at 300 kV equipped with a BioQuantum K3 Imaging Filter (Gatan) slit width of 20eV and a Falcon 4i electron detector with a Selectris X (ThermoFisher) slit width of 10eV. A total of 12,630 movies of the ADGRV1β/RE02 complex were acquired in counting mode using the EPU software (Thermo Fisher Scientific) (**Table S3**).

Image processing was performed using CryoSPARC (v4.0.3 or later) ^38^. Movies were motion-corrected using Patch Motion Correction, and contrast transfer function (CTF) parameters were estimated using Patch CTF estimation. Initial particle picking was performed with the Blob Picker tool, followed by several rounds of 2D classification. The 2D class averages displaying secondary structure elements served as templates for a second round of particle picking (1,826,101 particles). After additional rounds of 2D classification, classes showing high-resolution features, such as the transmembrane segments of ADGRV1β, and containing only one complex per detergent micelle were selected and extracted to generate a reference-free 3D ab initio model. Particles were further classified through iterative rounds of 3D classification (without alignment), heterogeneous refinement, and local refinement using static masks that excluded the detergent micelle surrounding the complex. These masks were necessary to resolve the GPCR region of ADGRV1β and the interface with RE02 at high resolution. A final reconstruction at 3.8 Å resolution was obtained with a set of 97, 447 particles.

### Model building and refinement

AlphaFold ^39^ models of ADGRV1β and the nanobody were manually fitted into the cryo-EM map. Following the removal of the unstructured N-terminal regions of ADGRV1β from the model, the complex was subjected to iterative cycles of manual rebuilding using Coot ^40^ and real-space refinement using Phenix ^41^.

## Data availability

The cryo-EM map and model coordinates of the ADGRV1β/RE02 complex have been deposited at the EMDB and at the PDB, respectively under accession codes 50743 and 9FTE.

## MD simulations setup, equilibration, production, and analysis

Atomistic MD simulations of ADGRV1β in explicit water and POPC bilayer were performed in the presence or absence of the VHH starting from the cryo-EM structure in inactive state. Initial models were built using MODELLER ^42^ v. 10.5 by adding to the cryo-EM structure the 14 N-terminal residues (residues 5884-5897) as well as other missing residues/atoms. The three top scoring models were selected with DOPE ^43^ and three additional models were generated after removing the VHH from each of the model. CHARMM-GUI ^44^was used to setup the systems for the MD simulations. Each of the six models was inserted in a homogeneous POPC lipid bilayer and solvated in a triclinic water box. K+ and Cl-were added to ensure charge neutrality at concentration equal to 0.15 M. Additional details of the MD setup are reported in **Table S5**. The CHARMM36m ^45^ forcefield was used for proteins and lipids and the mTIP3P ^46^ model for water molecules. The standard multi-step preparation protocol of CHARMM-GUI was used to energy-minize and gradually equilibrate the system at a temperature of 303 K and pressure of 1 atm using positional restraints on protein and lipid atoms. The last equilibration step with positional restraints applied only to protein backbone atoms was performed for 50 ns. In all production simulations the equations of motion were integrated by a leap-frog algorithm with timestep equal to 2 fs together with LINCS constraints on h-bonds ^47^. The smooth particle mesh Ewald method ^48^ was used to calculate electrostatic interactions with a cutoff equal to 1.2 nm. Van der Waals interactions were gradually switched off at 1.0 nm and cut off at 1.2 nm. Temperature was maintained at 303 K using the Bussi-Donadio-Parrinello thermostat ^49^. Pressure was maintained at 1 atm using a semi-isotropic Parrinello-Rahman barostat ^50^ to allow deformations in the *xy* plane independent from the *z*-axis. The six production simulations were run for 2µs each. Configurations were saved to the trajectory file every 10 ps. All simulations were carried out using GROMACS v. 2022.5 ^51^.

Analysis of the production simulations was carried out with MDAnalysis v. 2.7 ^52^. The stability of ADGRV1β in the presence or absence of the VHH was assessed by calculating the backbone Root Mean Square Deviation (RMSD) from the cryo-EM structure of ADGRV1β after excluding the VHH, the first 14 N-terminal residues, and the ICL3 loop (residues 6079-6095). The position of the ICL3 loop during the production simulations was assessed by computing the z-coordinate of the centre of mass of the ICL3 backbone atoms after aligning the centre of the lipid bilayer at z=0. To quantify the number of salt bridges formed by the ICL3 loop with the rest of ADGRV1β, for each frame of the trajectories we used MDAnalysis to monitor the distances between the sidechain charged groups of aspartic acids (OD1/OD2), glutamic acids (OE1/OE2), lysines (NZ), and arginines (NH1/NH2). A salt bridge was then defined as formed if the distance between groups with opposite charge was lower than 0.32 nm. To monitor the formation of hydrogen bonds, we used the *Hydrogen Bond Analysis* module of MDAnalysis. A donor-acceptor distance and angular cutoffs of 0.3 nm and 150° were used to define the formation of a hydrogen bond. To quantify the number of hydrophobic contacts between the ICL3 loop and the rest of ADGRV1β, we computed for each frame of the trajectory the distance between the Cα atoms of the hydrophobic residues (alanine, isoleucine, leucine, methionine, phenylalanine, tryptophan, tyrosine, and valine). A hydrophobic contact was then defined as formed if the distance between two Cα atoms was lower than 0.8 nm.

## Acknowledgments

The NanoImaging Core at Institut Pasteur is acknowledged for support with sample preparation, image acquisition. The NanoImaging Core was created with the help of a grant from the French Government’s Investissements d’Avenir program (EQUIPEX CACSICE -Centre d’analyse de systèmes complexes dans les environnements complexes, ANR-11-EQPX-0008). We gratefully acknowledge the support of the Swing Beamline (Synchrotron SOLEIL, Saint Aubin) and Aurélien Thureau for assistance with SAXS experiments. This work was supported by the Fondation pour l’Audition (program N°2019-0018C1) awarded to N.W., the Ministère de l’Enseignement Supérieur et de la Recherche (grant N°4133/2021) to Y. A. (grant N°3178/2018) to B.C-C, and the Fondation pour la Recherche Médicale (grant N°FDT202106013076) to B.C-C. We thank Nathalie Barilone (Institut Pasteur, France) for her support in the production of proteins and cell cultures. The authors acknowledge the Production and Purification of Recombinant Proteins core facility (Institut Pasteur, France) for the cell culture support, and the Arpège core facility (Montpellier, France) for functional assays of ADGRV1. We also thank Pierre-Jean Corringer (Institut Pasteur, France) and Guillaume Lebon (IGF Montpellier, France) for helpful discussions and advice.

