## Supplemental Figure 1 for "Structural insights into the inactive state of the adhesion GPCR ADGRV1"

Supplementary figures and tables

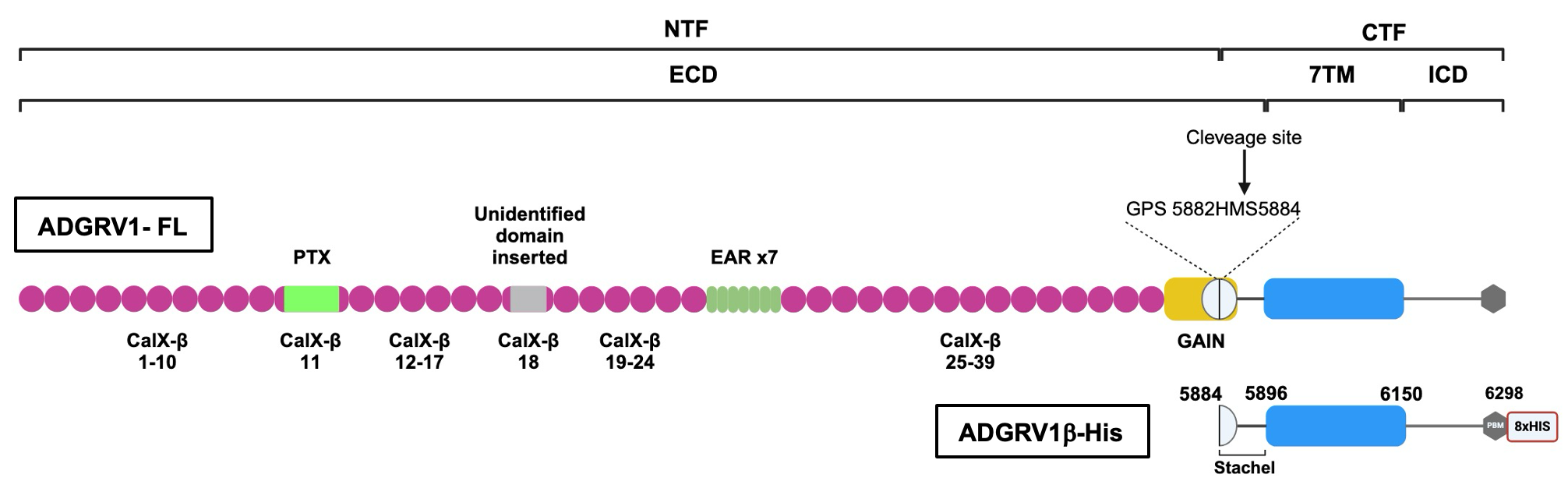

**Figure S1.** Schematic representation of structural organization of ADGRV1. ADGRV1β-His construct is used in our biophysical and cryo-EM studies. Indicated delimitations correspond to the mouse protein sequence.

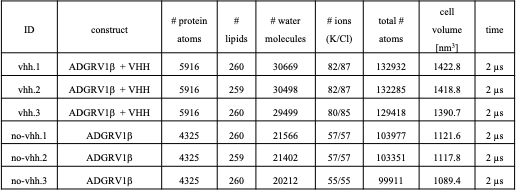
