## Supplemental Figure 2 for "Structural insights into the inactive state of the adhesion GPCR ADGRV1"

Supplementary figures and tables

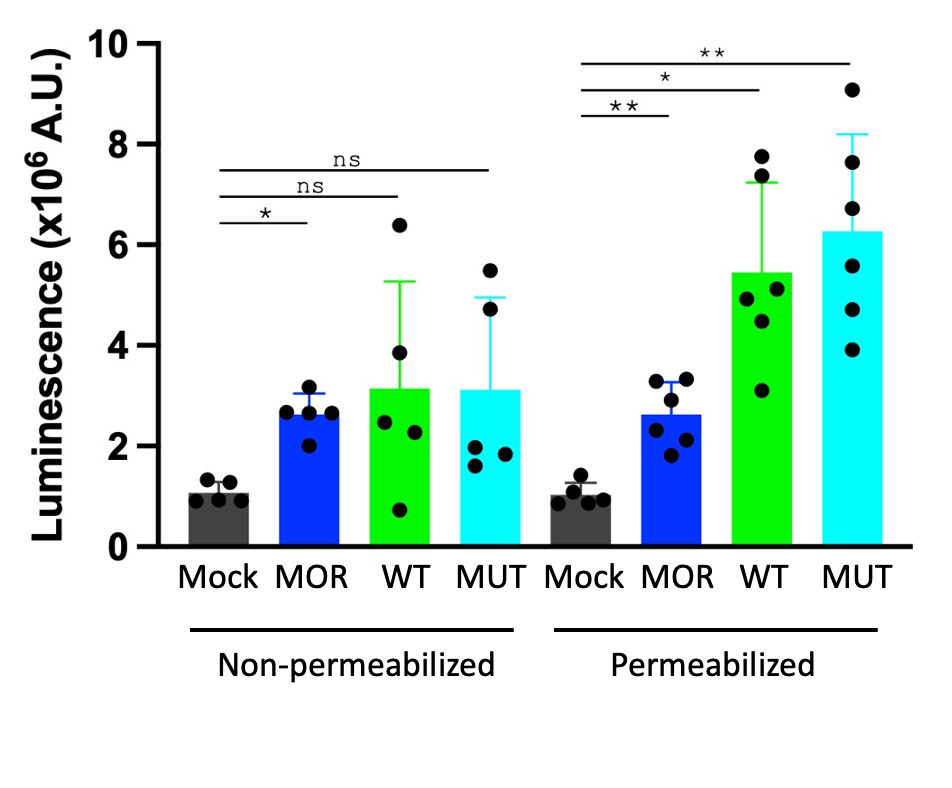

**Figure S2.** ADGRV1β WT and ADGRV1β MUT receptors expression monitored by ELISA assay on non-permeabilized or permeabilized HEK cells. Cells expressing MOR receptor are used as positive control and Mock cells (vehicle treated cells) as negative control.
