## Supplemental Figure 3 for "Structural insights into the inactive state of the adhesion GPCR ADGRV1"

Supplementary figures and tables

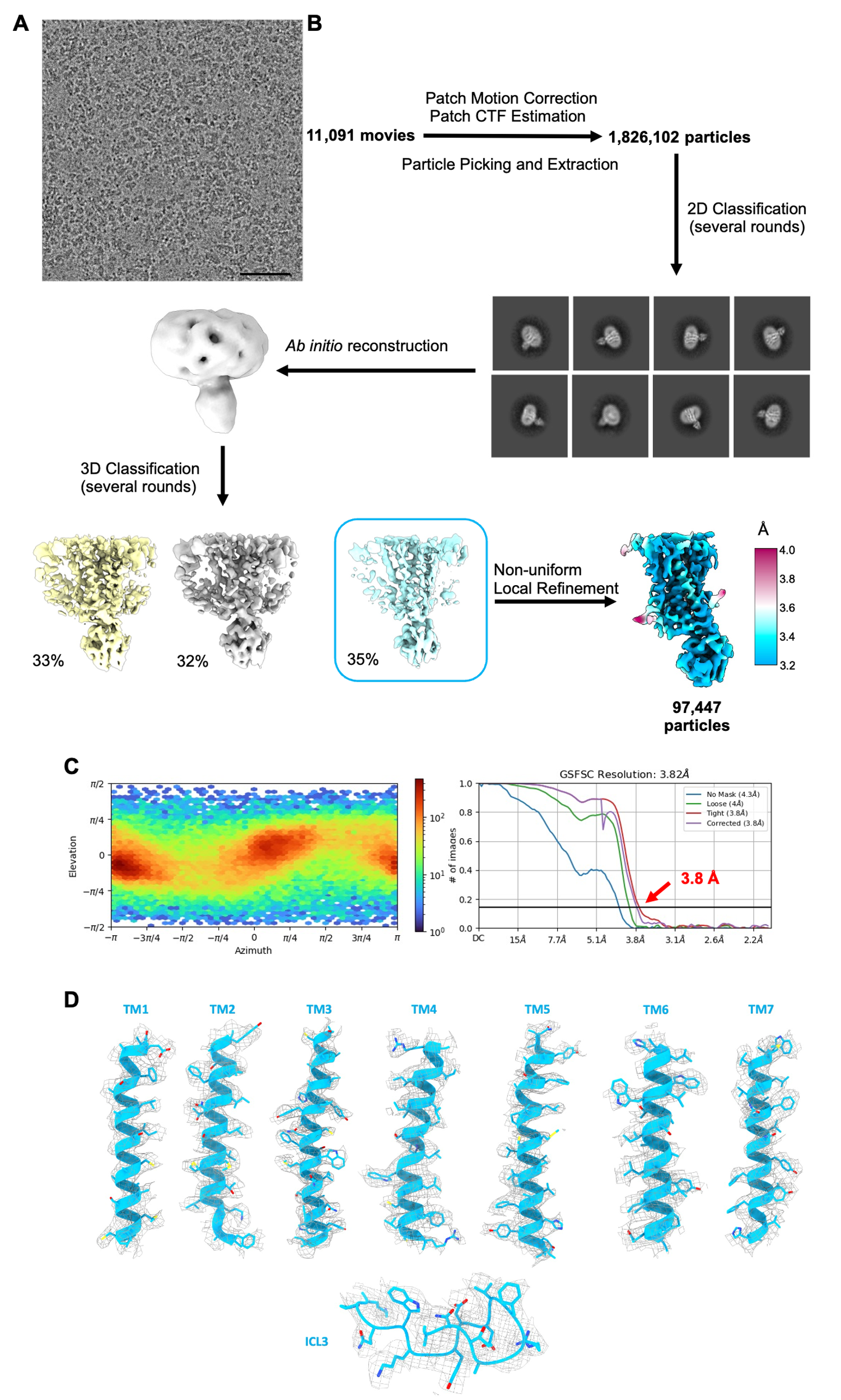

**Figure S3.** ***Cryo-EM data processing of inactive ADGRV1*β** ***and RE02 complex****, related to figure 3. A) Representative cryo-EM micrograph (scale bare: 60nm) of ADGRV1*β*-RE02 complex. B) Flow chart of cryo-EM data processing and cryo-EM maps of inactive ADGRV1*β*-RE02 complex colored by local resolution. C) Gold-standard FSC curve of the final refined map and angular distribution of particles used in the final 3D reconstitution the complex. D) Cryo-EM density maps and models are shown for the seven- transmembrane helices and the ICL3.*
