## Supplemental Figure 4 for "Structural insights into the inactive state of the adhesion GPCR ADGRV1"

Supplementary figures and tables

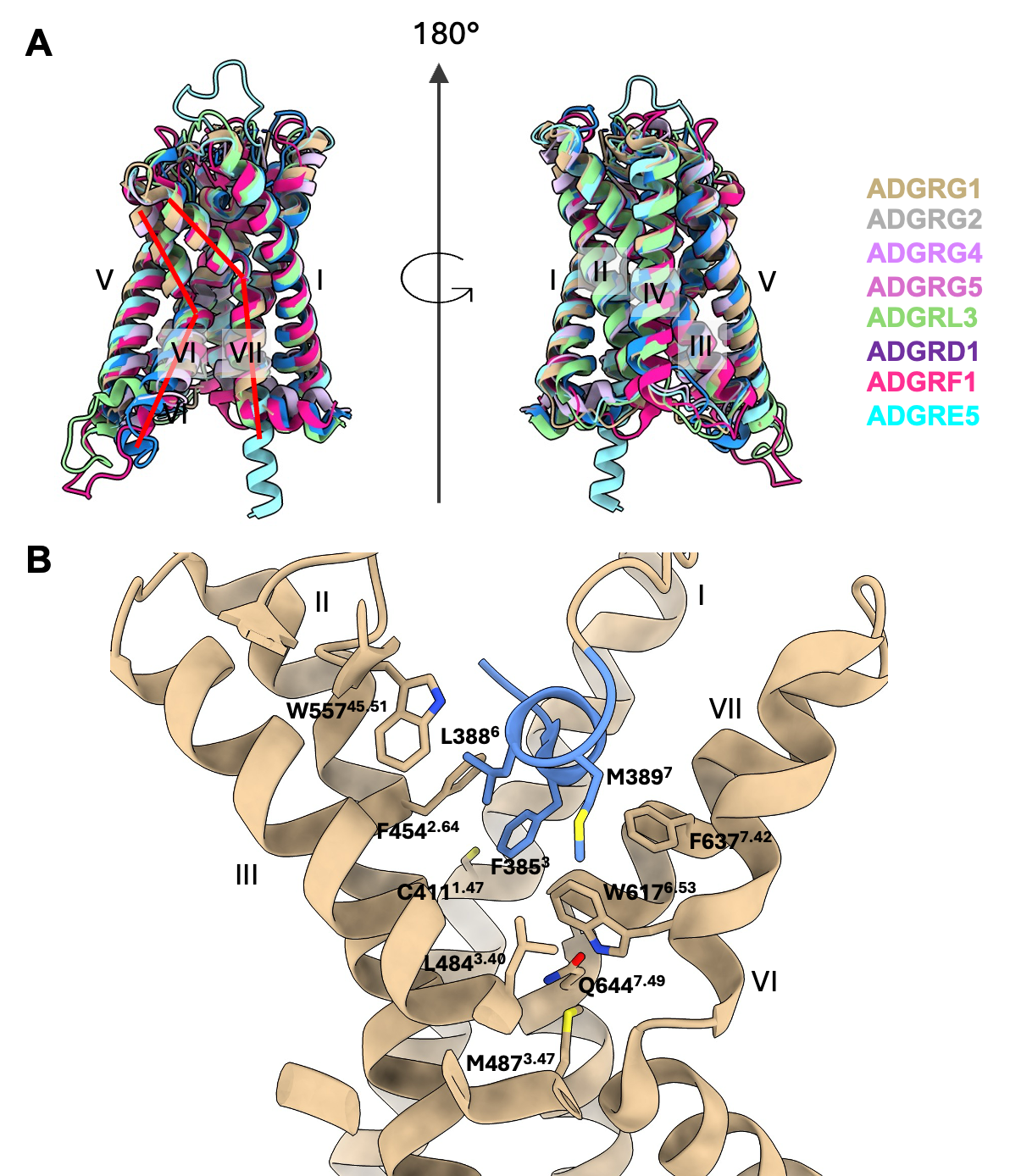

**Figure S4.** A) Superimposition of reported Stachel sequence-activated aGPCR structures. ADGRG1-G13 (PDB: 7SF8, ochre); ADGRG2-Gs (PDB:7WUQ, grey); ADGRG4-Gs (PDB:7WUJ, Lilac); ADGRG5- Gs (PDB: 7EQ1, light blue); ADGRL3-G13 (PDB: 7SF7, green); ADGRD1-Gs (PDB: 7WU2, grape); ADGRF1-Gi (PDB: 7WU4; Strawberry); ADGRE5-G13 (PDB: 8IKL, Blue). B) View of ligand pocket in the 7TM of ADGRG1 (ochre, PDB:7SF8) accommodating the TA Stachel peptide (Cornflower Blue).
