## Supplemental Figure 5 for "Structural insights into the inactive state of the adhesion GPCR ADGRV1"

Supplementary figures and tables

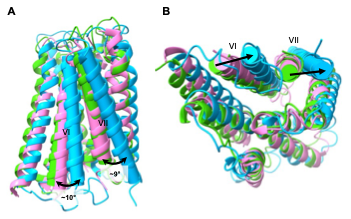

**Figure S5.** A) Superimposition of ADGRE5 (PDB: 8IKL) and ADGRL3 (PDB: 7SF7) structures with the structure of ADGRV1β in their inactive state, showing the shift of TMVI and TMVII in ADGRV1β. B) Extracellular view of TMVI and TMVII shift.
