## Supplemental Figure 6 for "Structural insights into the inactive state of the adhesion GPCR ADGRV1"

Supplementary figures and tables

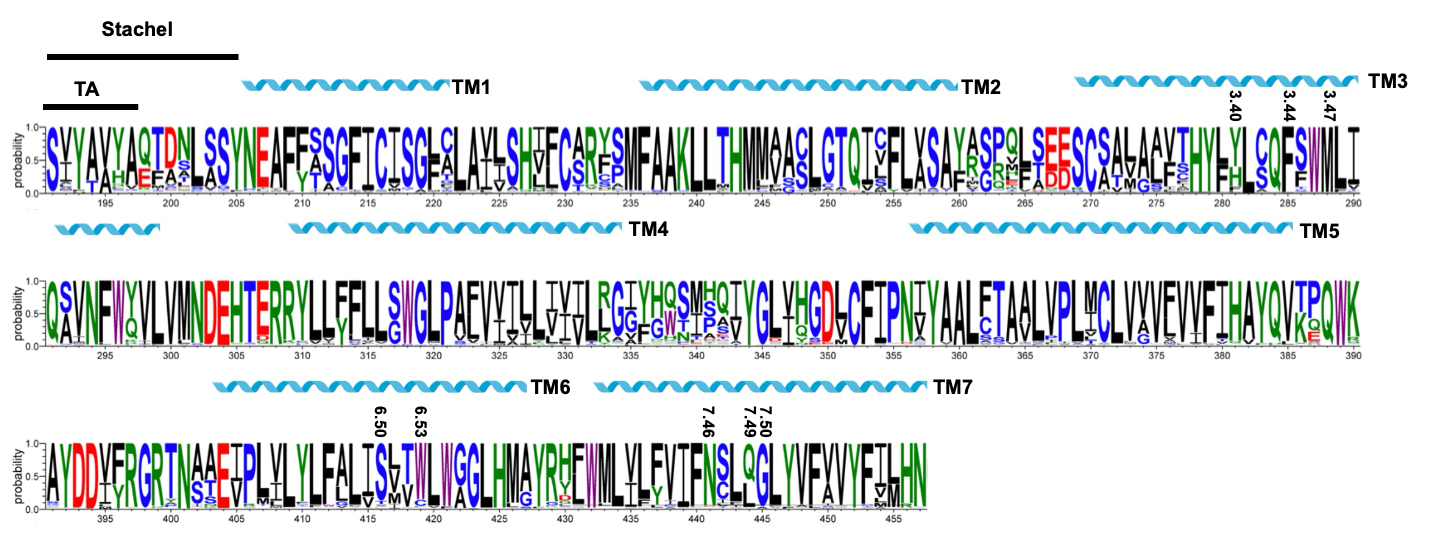

**Figure S6.** Logo representation of ADGRV1β sequences from 97 orthologs retrieved from the UniprotKB database using Hidden Markov Profile, with subsequent sequence length filtering. The N-terminal TA of the Stachel peptide, TM and key positions in the TM are highlighted.
