## Supplemental Figure 7 for "Structural insights into the inactive state of the adhesion GPCR ADGRV1"

Supplementary figures and tables

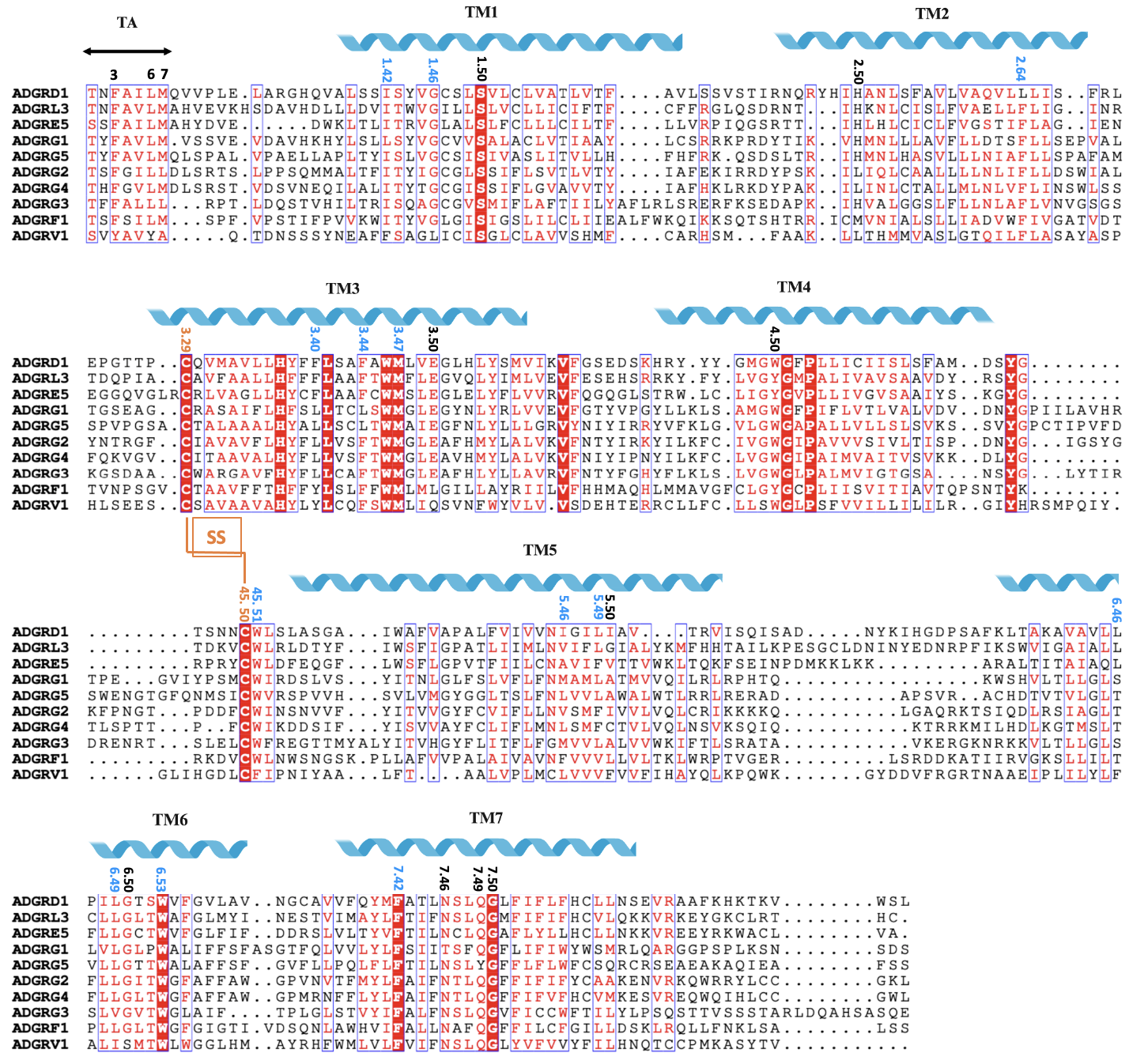

**Figure S7.** Sequence alignment of aGPCR resolved by cryo-EM and ADGRV1β. The N-terminal TA of the Stachel peptide, TM and key positions in the TM are highlighted. The alignment was generated using MAFFT and the graphic was prepared on the ESPript3.0 server.
