## Supplemental Table 1 for "Structural insights into the inactive state of the adhesion GPCR ADGRV1"

Supplementary figures and tables

**Table S1.** AUC data collection and derived parameters.

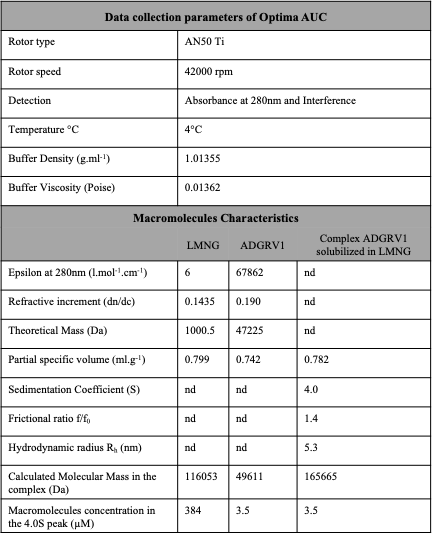
