## Supplemental Table 2 for "Structural insights into the inactive state of the adhesion GPCR ADGRV1"

Supplementary figures and tables

**Table S2.** SAXS data collection and scattering derived parameters.

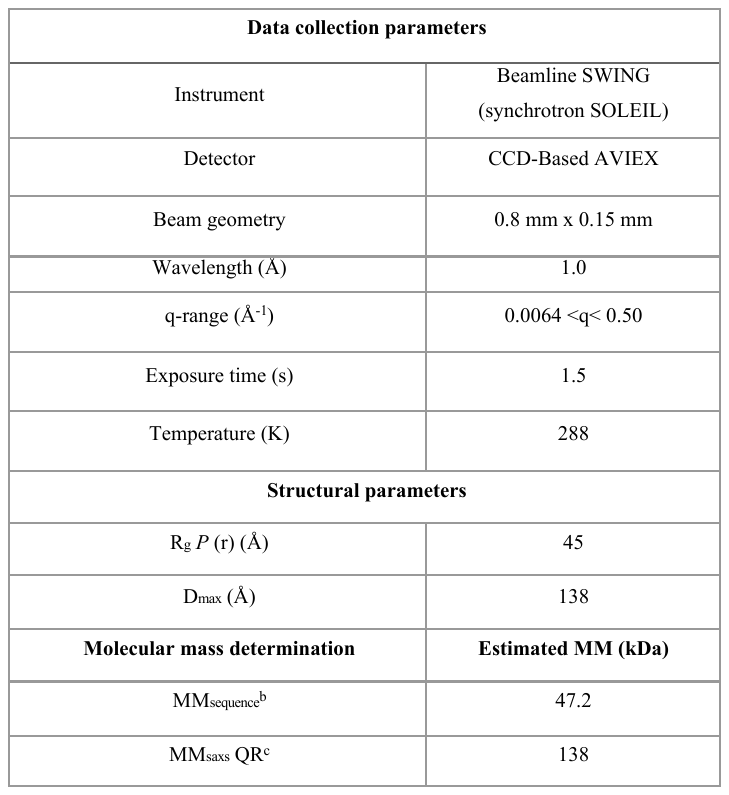
