## Supplemental Table 3 for "Structural insights into the inactive state of the adhesion GPCR ADGRV1"

Supplementary figures and tables

**Table S3.** Cryo-EM data collection, model refinement and validation statistics.

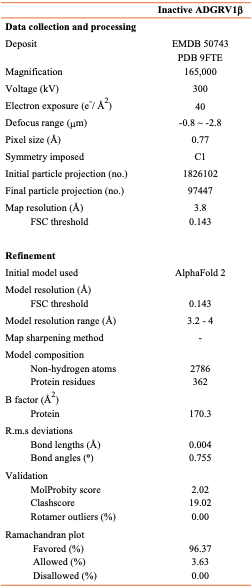
