## Supplemental Table 4 for "Structural insights into the inactive state of the adhesion GPCR ADGRV1"

Supplementary figures and tables

**Table S4.** List of the residues without side chains assignements within the map density**.**

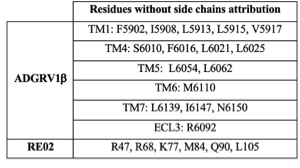
