## Supplemental Table 5 for "Structural insights into the inactive state of the adhesion GPCR ADGRV1"

Supplementary figures and tables

**Table S5.** **Details of the MD simulations**. The table reports: simulation ID, construct, number of protein atoms, number of POPC lipid molecules, number of water molecules, number of K and Cl ions, total number of atoms, volume of the simulation cell, and total simulation time.

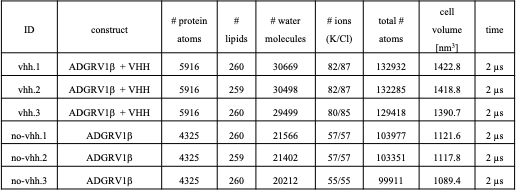
